# PCNA is a Nucleotide Exchange Factor for the Clamp Loader ATPase Complex

**DOI:** 10.1101/2025.07.02.662830

**Authors:** Joshua Pajak, Jacob T. Landeck, Xingchen Liu, Krishna Anand, Sasha Litvak, Brian A. Kelch

## Abstract

All life requires loading ring-shaped sliding clamp protein complexes onto DNA. The sliding clamp loader is a conserved AAA+ ATPase that binds the sliding clamp, opens the ring, and places it onto DNA. While recent structural work on both the canonical and ‘alternative’ clamp loaders has shed light into how these machines perform their task once, it remains unclear how clamp loaders are recycled to load multiple sliding clamps. Here, we present structures of the *Saccharomyces cerevisiae* clamp loader Replication Factor C (RFC) in absence of sliding clamp or supplemented nucleotide. Our structures indicate that RFC holds onto ADP tightly in at least two of its four ATPase active sites, suggesting that nucleotide exchange is regulated. Our molecular dynamics simulations and biochemical data indicate that binding of the sliding clamp PCNA causes rapid exchange of tightly bound ADP. Our data suggests that PCNA acts as a nucleotide exchange factor by prying apart adjacent subunits, providing a pathway for ADP release. We propose that, by using its own substrate as a nucleotide exchange factor, RFC excludes off-pathway states that would arise from binding DNA prior to PCNA.

## Introduction

Sliding clamps are protein complexes that form closed rings around DNA and act as scaffolds to coordinate the function of over two hundred protein partners (1–3). These partners, such as DNA polymerases and endonucleases, act on DNA during replication or repair. Thus, sliding clamps are crucial for maintaining genomic integrity. The eukaryotic sliding clamp is <u>P</u>roliferating <u>C</u>ell <u>N</u>uclear <u>A</u>ntigen (PCNA), a homo-trimeric protein complex. Because PCNA is a closed ring in solution, it cannot encircle DNA on its own. Instead, it is loaded by a clamp loader complex.

The canonical clamp loader in eukaryotes is <u>R</u>eplication <u>F</u>actor <u>C</u> (RFC) (1). RFC is a hetero-pentameric <u>A</u>TPase <u>A</u>ssociated with diverse cellular <u>A</u>ctivities (AAA+) protein complex (4, 5). The five subunits of RFC are named A through E, and each contain a AAA+ module. The AAA+ module typically binds ATP in the cleft formed between the Rossmann domain and the lid subdomain, and each ATPase active site is completed by an arginine finger residue that is donated *in trans* by a neighboring subunit. RFC uses the energy of ATP binding to open and load PCNA onto DNA, and ATP hydrolysis and/or phosphate release trigger dissociation of RFC from PCNA that is wrapped around DNA (1, 6).

During lagging strand synthesis, RFC loads PCNA onto each Okazaki fragment (7, 8). Because Okazaki fragments are approximately 165 bp in length in yeast (9), each replication cycle requires thousands of PCNA loading events. Thus, once RFC hydrolyzes ATP and dissociates from the loaded PCNA, it must rapidly exchange its spent ADP for new ATP to begin the process anew. *In vitro* steady-state and rapid kinetics characterization have demonstrated that RFC loads PCNA at a rate of ∼1 s^-1^ and that dissociation of RFC from loaded PCNA is the rate-limiting step (6, 10, 11). Thus, while the rate of nucleotide exchange has not yet been measured, it is expected to be relatively fast, as it is not rate-limiting.

Here, we show that RFC counter-intuitively binds ADP tightly in the absence of PCNA, with a dissociation rate that is orders of magnitude slower than the rate of clamp loading. To overcome this kinetic barrier, we find that PCNA acts as a nucleotide exchange factor. The additional regulation of using its substrate as an ATP exchange factor likely helps ensure that RFC binds PCNA prior to DNA and gives rise to a well-ordered loading reaction that avoids futile cycles.

## Results

### The A subunit’s AAA+ domain is flexibly tethered to the core RFC complex

To probe how RFC exchanges ADP for ATP, we sought to determine the structure of the apo state of RFC. Toward this goal, we determined the structure of recombinant yeast RFC with no added substrate (i.e. no nucleotide, PCNA, or DNA) using single-particle cryogenic electron microscopy (cryo-EM). Our data processing resulted in six 3D classes ranging in overall resolution between 2.5 and 2.7 Å gold-standard Fourier Shell Correlation (FSC) (**Fig. 1A, Fig. 1S1-1S5**), allowing us to unambiguously model backbone and sidechains within the maps.

**Figure 1.**
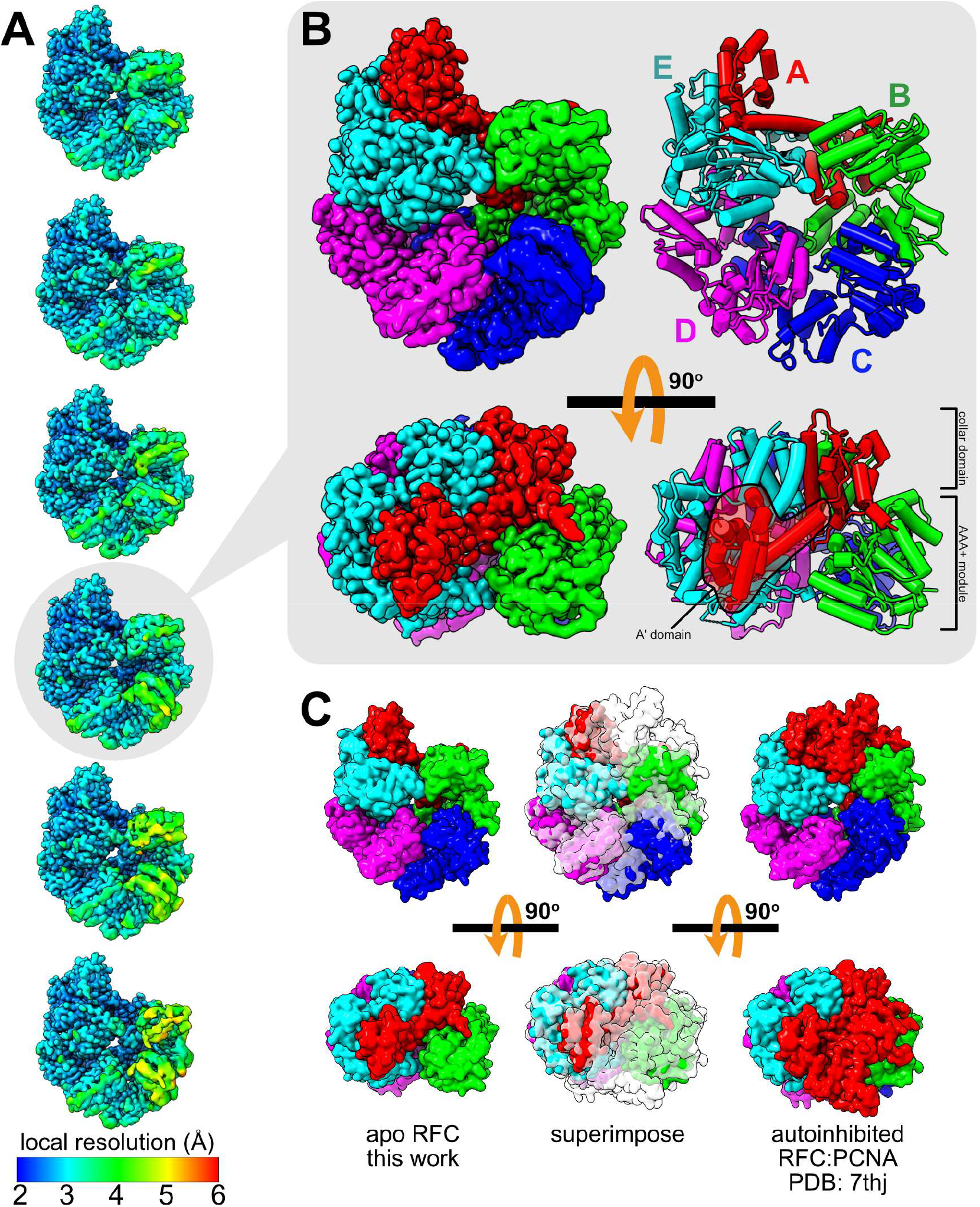
Cryo-EM reconstructions of the apo yeast RFC complex. **A**. Six reconstructions after 3D classification are depicted as viewed from the PCNA-interacting interface, and color coded by local resolution estimation. **B**. A single reconstruction is highlighted. (left) The reconstruction and model are is color coded by the RFC subunit (right). **C**. Gaussian surfaces of the model shown in panel B (left) and the autoinhibited RFC:PCNA complex (right; PDB: 7thj; PCNA not shown for clarity) are superimposed (center; apo colorful, autoinhibited transparent) to highlight the missing density corresponding to the AAA+ domain of the A subunit.

In all six classes, we observe no density corresponding to the AAA+ domain of the A subunit, despite strong density for the collar and A′ domain (**Fig. 1B,C**). Density suficient for building a model of the A subunit does not appear until residue ∼544, although diffuse density extending towards the N-terminus does appear at lower contours (**Fig. 1S5**). This helical motif of the A subunit (residues 543-547) unwinds as RFC opens PCNA (12), demonstrating its plasticity. Thus, our reconstructions suggest that the AAA+ module of the A subunit is flexibly tethered to the core of the RFC complex. Recent studies of the alternative clamp loader CTF18-RFC from both humans and yeast have shown that the AAA+ domain of CTF18 (which replaces Rfc1 as the A subunit) is also flexibly tethered to the core RFC complex (13–15). Thus, loose tethering of the A subunit may be a common feature of eukaryotic clamp loaders and may be related to how the clamp loader dissociates from the sliding clamp after ATP hydrolysis and phosphate release.

### The remaining AAA+ domains have mixed nucleotide occupancy and varying degrees of flexibility

In all six of our 3D classes, we observe the AAA+ modules of the B, C, D, and E subunits (**Fig. 1B,C**). Inspection of the ATPase active sites in our reconstructions reveals bound nucleotide. We observe GDP bound to the E subunit in all six 3D classes (**Fig. 2A**), in agreement with previous high-resolution reconstructions of RFC:PCNA that also identified GDP in this catalytically inactive site (16). As in Schrecker & Castaneda *et al*., we did not introduce GTP or GDP at any step during protein purification or grid sample preparation, so the bound GDP is co-purified with RFC. This finding is not entirely surprising as GDP likely plays a structural role to stabilize the E subunit (17).

**Figure 2.**
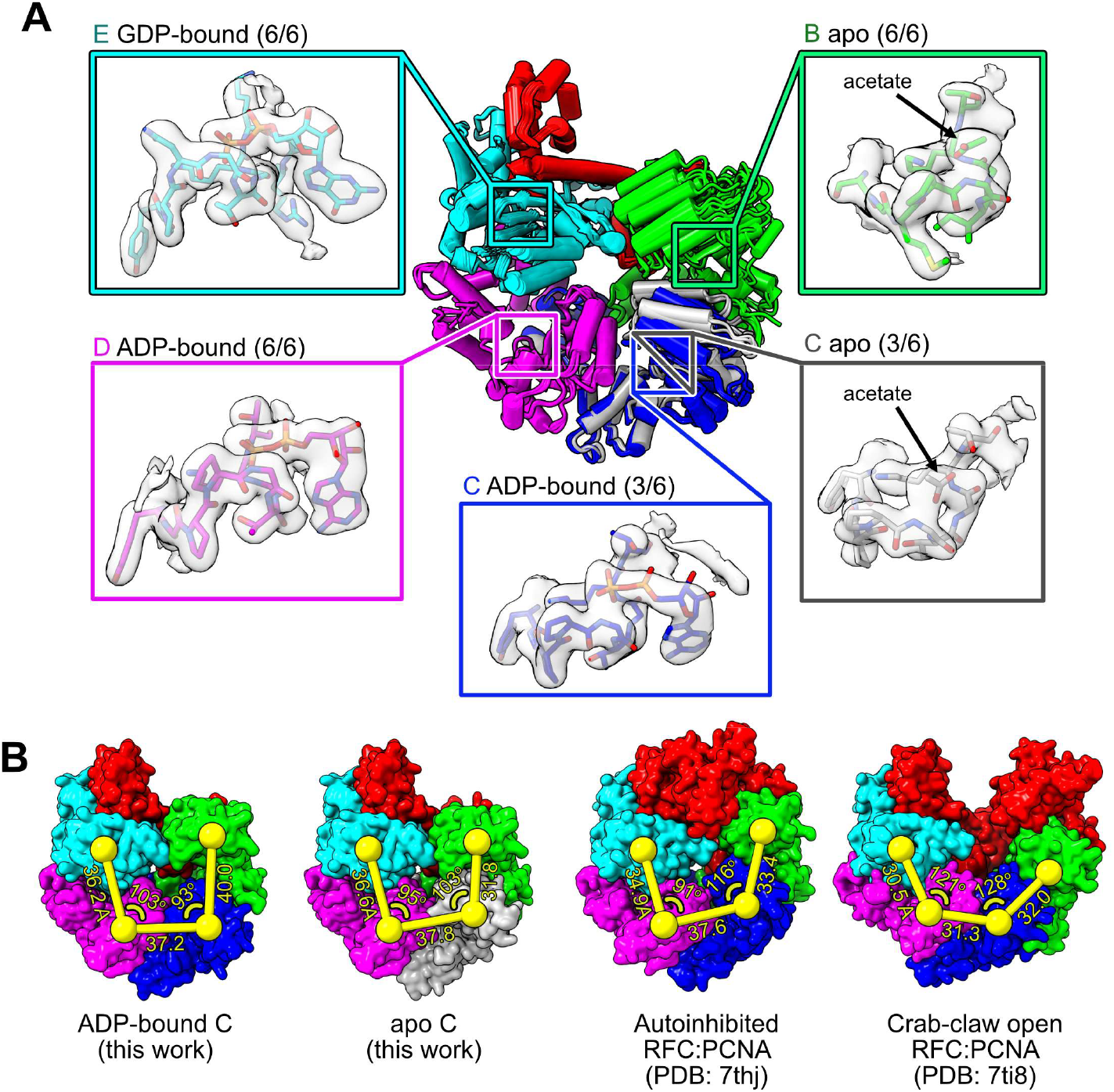
Cryo-EM reconstructions reveal mixed nucleotide occupancy of the ‘apo’ RFC complex. **A**. All six models are superimposed on the D subunit. The C subunit is silver if it is apo and blue if it is ADP-bound. Sub-panels show the Walker A motif and bound nucleotide from a representative class. **B**. Centers-of-mass of the Rossmann domains of the A-D subunits are shown as yellow spheres and superimposed on Gaussian surfaces of modeled RFC. Two models from this work are shown, as well as the autoinhibited RFC:PCNA complex (PDB: 7thj; PCNA not shown for clarity) and the crab-claw open RFC:PCNA complex (PDB: 7ti8; PCNA not shown for clarity).

Surprisingly, we also observe density corresponding to ADP in the active site of the D subunit in all six 3D classes and in the active site of the C subunit in three of six 3D classes (**Fig. 2A**). There is density attributable to the adenine ring, ribose sugar, and α- and β-phosphates, but there is no density attributable to a γ-phosphate, allowing us to rule out that ATP is bound at these sites. Unlike the E subunit, the D and C subunits are active ATPase sites and ATP hydrolysis at these sites is required for eficient clamp loading (17, 18). Because RFC loads PCNA with a maximal velocity of 1 s^-1^ (6, 10), one might expect that these sites release ADP rapidly. However, as neither ATP nor ADP was added to this sample, ADP must also have co-purified with RFC. Thus, ADP is released slowly. This puzzling observation implies that nucleotide exchange in RFC is catalyzed by some component not included in our cryo-EM sample preparation.

We observe no density attributable to nucleotide in the active site of the C subunit in the remaining three 3D classes nor in the active site of the B subunit in any of the six 3D classes (**Fig. 2A**). While there is density directly abutting the Walker A motif in these active sites, this density cannot account for the adenine ring or ribose sugar. Due to the size of this density, we assign it to be an acetate ion, which was included in our buffer conditions; acetate can act in lieu of a phosphate and coordinate the P-loop backbone nitrogens.

When aligned to the D subunit, our structures exhibit flexibility in the relative positions of the B and C subunits, while the E subunit and A′ domain remain rigid (**Fig. 2A and Movie S1**). The AAA+ domain of the C subunit assumes one of two conformations depending upon nucleotide occupancy (compare blue and silver subunits in **Fig. 2A**).

The overall arrangement of AAA+ domains is unique from those previously observed when RFC is ATP-bound and contacting PCNA (**Fig. 2B**). Similar to the autoinhibited state, the AAA+ domains form a tight spiral, and subunits are generally far from one another compared to the crab-claw open conformation. This arrangement prevents completion of the active site by catalytic residues that are donated *in trans* (12).

### Nucleotide binding/release and ATP hydrolysis rotate the lid subdomains of RFC subunits

Despite extensive structural characterization of eukaryotic clamp loader complexes (12–16, 19–28), all but two of these structures were determined in the presence of a slowly-hydrolysable ATP analog (ATP-γS). The exceptions are an ATP-bound alternative clamp loader (CTF18-RFC) (15) and an ADP-bound alternative clamp loader (Rad24-RFC) (21). Hence, it has not been possible to determine the structural changes associated with ATP hydrolysis and phosphate release in the canonical RFC complex, or nucleotide binding/release in any clamp loader complex. The mixed nucleotide occupancy of our reconstructions allows us to outline some of these structural changes for the first time.

To understand the conformational changes induced by ATP binding, we aligned ATP-γS bound B and C subunits from the autoinhibited RFC:PCNA complex (PDB: 7thj) to the apo B and C subunits (**Fig. 3A**). In both subunits, the lid subdomain rotates relative to the Rossmann fold, but the rotations are distinct in both magnitude and direction. When viewed from the subunit donating catalytic residues *in trans*, the lid subdomain of the C subunit rotates ∼27o clockwise while the lid subdomain of the B subunit rotates ∼15o counterclockwise. In many AAA+ ATPases, the motion of the lid subdomain is similar to that seen in the C subunit (29, 30). To our knowledge, this is the first time that a lid subdomain is observed rotating in the opposite direction in response to ATP binding. We note that in the autoinhibited structure, both the B and C subunits engage PCNA, which could affect this comparison. However, we see a similar rotation when comparing ADP-bound and apo C subunits, neither of which contact PCNA. The relatively large rotation of the C subunit in response to nucleotide binding supports the prior characterization of the C subunit as a ‘central swivel point’ (31).

**Figure 3.**
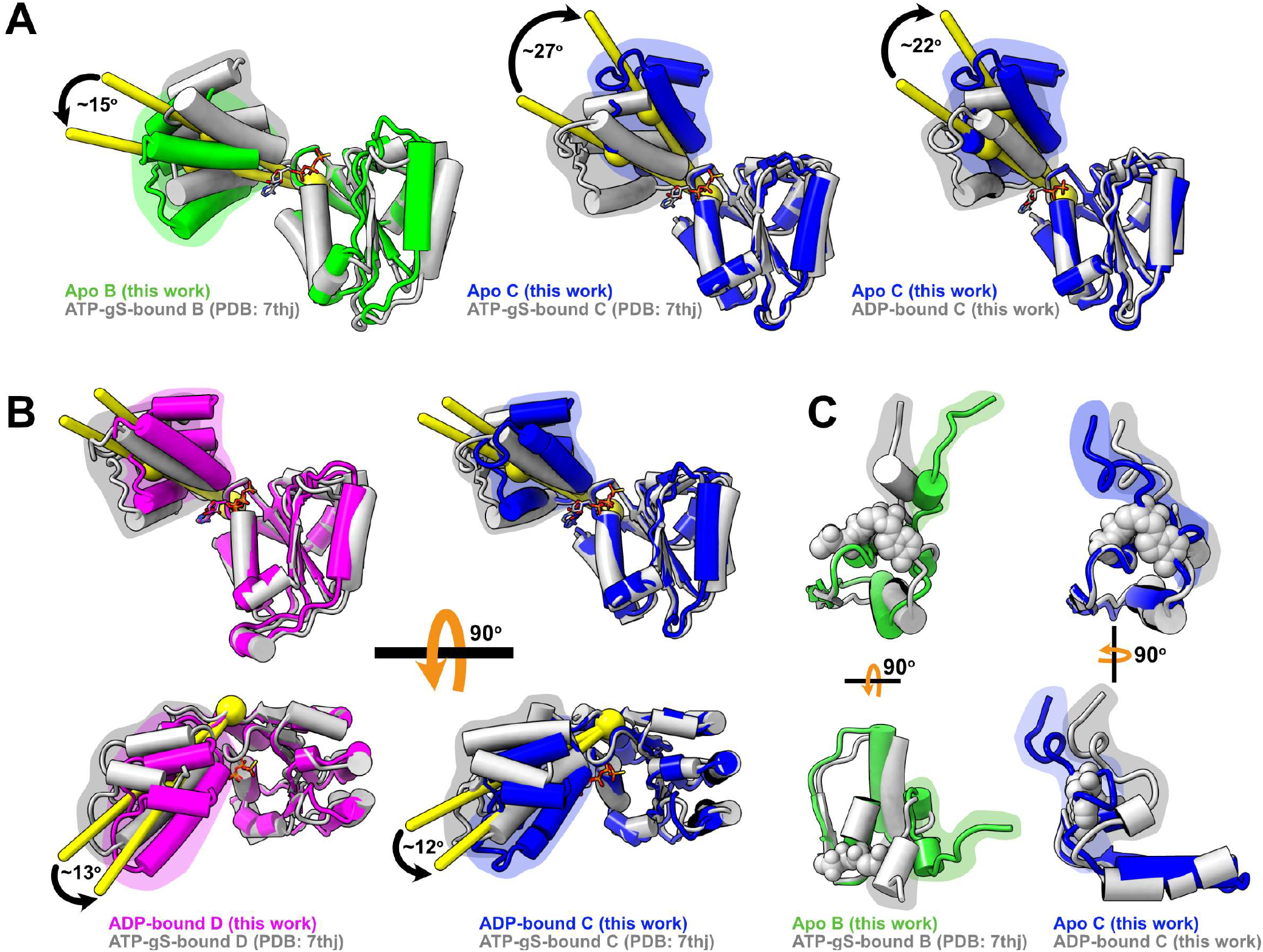
Nucleotide-dependent conformational changes of RFC subunits. **A**. Nucleotide-bound RFC subunits (ATP-γS-bound from the autoinhibited RFC:PCNA complex or ADP-bound from this work; silver) and apo subunits (colorful) are aligned on their ATPase active sites. The centers-of-mass of lid subdomains and the pivot point residues are shown as yellow spheres, and the angular displacement of the centers-of-mass is reported. The lid subdomain is highlighted by a shaded region. **B**. ADP-bound (colorful) and ATP-γS-bound (silver; from autoinhibited RFC:PCNA complex) subunits are aligned on their ATPase active sites. The centers-of-mass of lid subdomains and the pivot points are shown as yellow spheres, and the angular displacement of the centers-of-mass is shown. The lid subdomain is highlighted by a shaded region. **C**. Apo (colorful) and nucleotide-bound (silver) active sites are aligned. The N-terminal loop is highlighted by a shaded region.

Our structures also allow us to observe the conformational changes driven by ATP hydrolysis and phosphate release. We aligned our ADP-bound subunits to ATP-γS-bound subunits from the autoinhibited state. In both the C and D subunits there is a ∼12o rotation in the same direction, indicating that hydrolysis and phosphate release affect these subunits similarly (**Fig. 3B**). As in the above comparison, the ATP-γS bound C subunit contacts PCNA, which may complicate the interpretation. However, the D subunit does not contact PCNA in either of the two compared structures.

### The N-terminal loop can sterically block nucleotide

Each RFC subunit has an N-terminal loop that abuts the ATPase active site near the adenine ring and ribose sugar of ATP/ADP. In the nucleotide-bound conformation, this loop hydrogen bonds with the ribose sugar and adenine (**Fig. 3S1**). This loop extends past the active site and interacts with the lid subdomain. Because the B and C subunits lid subdomains rotate in opposite directions upon nucleotide release, their N-terminal loops are also pulled in opposite directions. In the apo B subunit, the N-terminal loop is pulled away from the active site, opening it for ADP release or ATP binding (**Fig. 3C**). However, the N-terminal loop of the apo C subunit occupies the active site, sterically blocking nucleotide binding (**Fig. 3C**). Thus, ATP binding may be not only prevented by slow ADP release but also by steric hindrance of apo subunits.

### Molecular dynamics simulations

#### predict fast ADP release from the A and B subunits and slow ADP release from the C and D subunits

To understand how ATP hydrolysis can affect the subunits’ dynamics, we performed all-atom molecular dynamics simulations of RFC bound with either ATP or ADP (except the E subunit, which was always GDP-bound). We performed three 2-μs long simulations of both nucleotide binding configurations. In general, the centers-of-mass of AAA+ domains are farther separated when ADP-bound compared to ATP-bound (**Fig. 4A**). This result indicates that the γ-phosphate of ATP helps bridge inter-subunit interactions. In particular, we find that the B and C subunits separate the most, potentially explaining why the B subunit does not retain ADP in our cryo-EM reconstructions.

**Figure 4.**
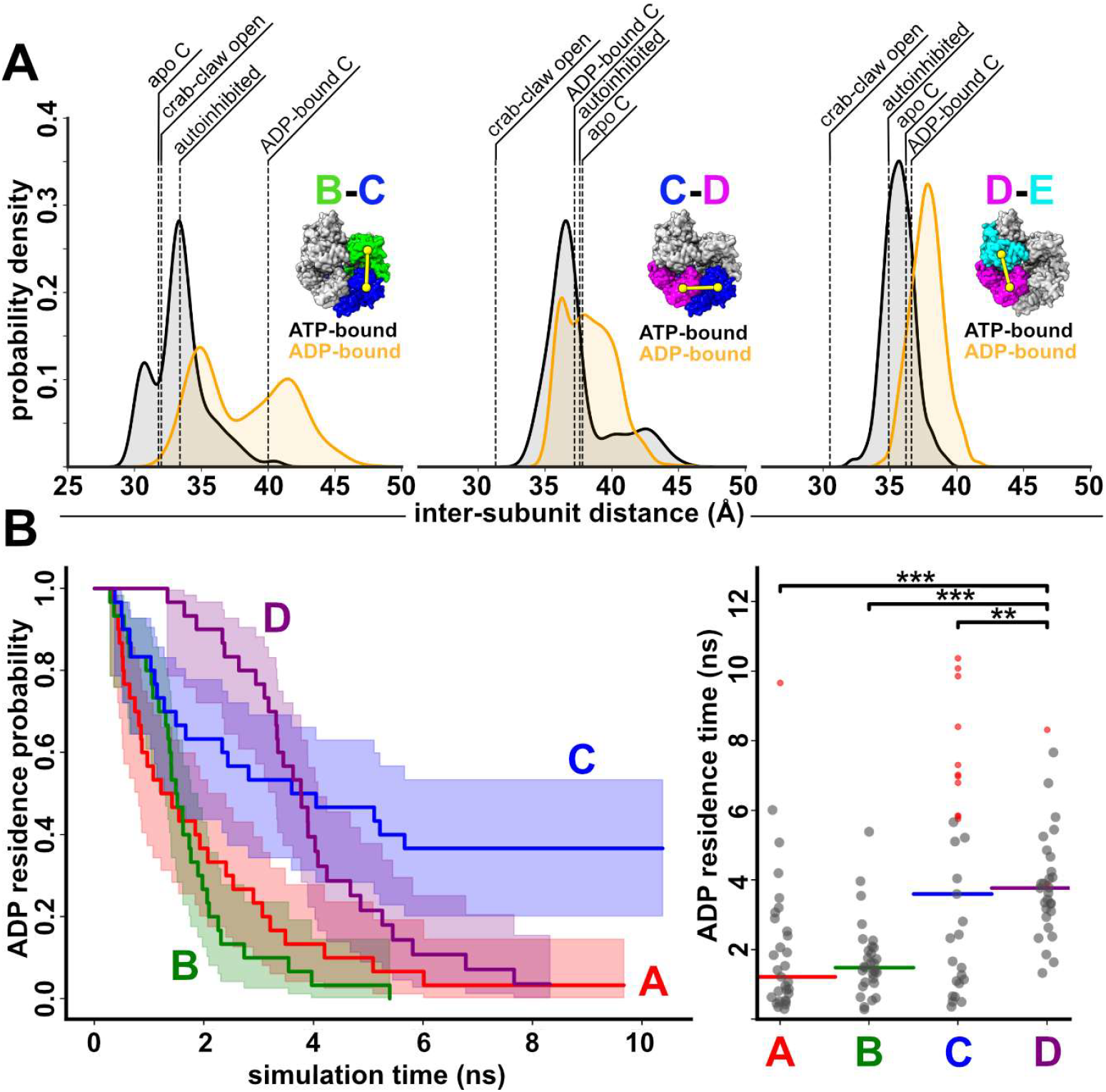
Molecular dynamics simulations of RFC and predicted relative ADP dissociation rates. **A**. The sampled distances between centers-of-mass of Rossmann domains from ATP-bound (black) and ADP-bound (orange) equilibrium molecular dynamics simulations are shown as kernel density estimates. The distances from the structures reported in **Fig. 2B** are shown as dashed lines for reference. **B**. Results from τ-RAMD simulations. (left) ADP residence probability is plotted against simulation time, and ADP residence time from each simulation is plotted on the right. Solid bars indicate the half-survival time, and red dots indicate right-censored data (where the unbinding event was not sampled during the simulation after one day of wall-time).

To test if the increased distance between subunits could give rise to different ADP release rates, we performed τ-Random Acceleration Molecular Dynamics (τ-RAMD) simulations (32). In these simulations, a biasing force is applied to ADP, which promotes dissociation from RFC in timescales that are computationally feasible. So, while the calculated dissociation times are not physically realistic, the relative rates are useful for comparison.

We seeded our τ-RAMD simulations from the endpoints of our three equilibrium MD simulations of ADP-bound RFC. From each of these seeds, we performed ten simulations for each ADP binding site from the A, B, C, and D subunits, for a total of 30 simulations per active site, or 120 total simulations. We measured the residence time of ADP in each simulation, and calculated the amount of time for half of the release events to occur (t_1/2_). The t_1/2_ of A and B are approximately ∼1.3 ns, and of C and D are approximately ∼3.7 ns. Thus, our τ-RAMD simulations predict that the A and B subunits release ADP faster than the C and D subunits (**Fig. 4B**). This result agrees with our cryo-EM reconstructions that show ADP in C and D but not in B.

#### *In vitro* characterization of ATP binding shows that PCNA accelerates nucleotide exchange

Our results thus far indicate that nucleotide exchange in RFC is catalyzed by some component not included in our cryo-EM sample preparation. There are a few possibilities for what that component may be. It could be binding PCNA or DNA, which were absent from our cryo-EM sample and MD simulations. This hypothesis is supported by previous data which show that RFC binds additional ATP molecules in the presence of PCNA or DNA (10, 33). Alternatively, RFC may use an interlaced exchange mechanism, whereby ATP binding at one subunit promotes ADP release and ATP binding at the neighboring subunit. Because our cryo-EM sample did not include ATP to start such an exchange mechanism, it may be possible that some release was permitted but the exchange cycle could not propagate around the AAA+ spiral.

To test these possibilities, we measured the association of a fluorescently labeled ATP analog 2’-(or-3’)-O-(N-Methylanthraniloyl)-ATP (MANT-ATP) to RFC. MANT-ATP binding to RFC was measured by Förster Resonance Energy Transfer (FRET), by using tryptophan residues proximal to the ATPase active sites as the donors (λ_EX_ = 297 nm) and MANT-ATP as the acceptor (λ_EM_ = 440 nm) (**Fig. 5A & Fig. 5S1**). Addition of RFC to MANT-ATP causes a FRET increase, but PCNA or p/t-DNA do not (**Fig. 5S2)**. We then monitored MANT-ATP fluorescence over time to observe exchange kinetics (**Fig. 5B & Fig. 5S3**). Upon addition of RFC, we observe an increase (∼1.23x over baseline) in MANT-ATP fluorescence within the dead time of mixing (∼20-30 seconds), indicating rapid MANT-ATP binding to RFC. Additionally, we observe a slow increase (from 1.23x to 1.29x over baseline) in MANT-ATP fluorescence over the course of hours as nucleotide continues to bind to RFC (**Fig. 5B**). The signal can be quenched by addition of excess unlabeled ATP at the end of the time course (1.29x to 1.07x) (**Fig. 5B & Fig. 5S2**). Taken together, these data indicate that ATP binds to some RFC subunits rapidly (within the dead time of the experiment), but to at least one subunit slowly.

**Figure 5.**
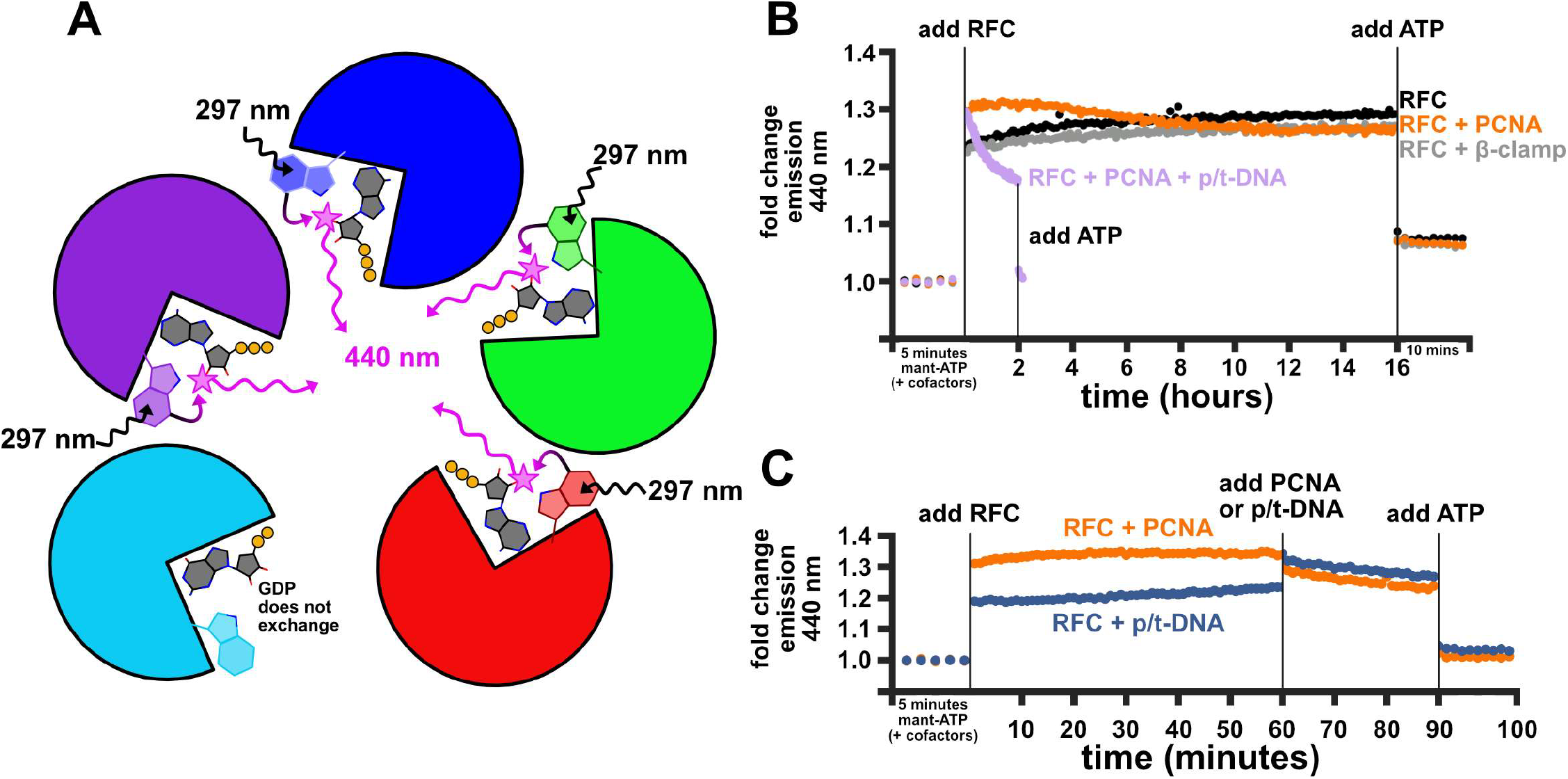
MANT-ATP FRET assays show that PCNA promotes fast nucleotide exchange in RFC. **A**. A cartoon schematic of the assay. Tryptophan residues are excited at 297 nm and can transfer energy if MANT-labeled ATP is bound, which emits at 440 nm. Because the E subunit is always GDP-bound, we assume that it does not exchange and contribute to signal in this assay. **B**. The fold change of measured emission at 440 nm is reported for different conditions. In all cases, the average intensity from the five minutes of measurement prior to adding RFC was used to set the baseline. **C**. Sequentially adding RFC’s binding partners shows that PCNA and not p/t-DNA is the primary driver of nucleotide exchange.

When MANT-ATP was preincubated with saturating PCNA, addition of RFC induces a large increase in FRET during the dead time (∼1.30x over baseline), followed by a plateau and slow decrease of fluorescence over the course of hours (from 1.31x to 1.26x) (**Fig. 5B**). We attribute the decrease in fluorescence to MANT-ATP hydrolysis. In support of this interpretation, we observe accelerated decay of FRET signal when MANT-ATP was preincubated with both saturating PCNA and p/t-DNA (conditions which maximize RFC’s ATPase activity; **Fig. 5B**). This behavior was not seen when MANT-ATP was preincubated with the sliding clamp from *E. coli*, which has low sequence similarity with PCNA (34) and should not bind RFC. This result illustrates that the rapid exchange is specific to PCNA.

To determine whether p/t-DNA can also promote nucleotide exchange, we preincubated with either PCNA or p/t-DNA, added RFC, and then later added in the missing component (**Fig. 5C & Fig. 5S3**). We observed that when pre-incubating MANT-ATP with p/t-DNA, addition of RFC only leads to a moderate increase in fluorescence within the dead time (∼1.19x over baseline, similar to that of RFC alone), followed by a moderate increase in fluorescence intensity over the course of one hour (from ∼1.19x to ∼1.24x). Adding PCNA to this sample rapidly increased fluorescence during the dead time of mixing (to ∼1.32x), followed by a decay in signal indicative of MANT-ATP hydrolysis (from 1.32x to 1.27x over 30 minutes). However, when the addition of p/t-DNA and PCNA were reversed, we observed a different trend. We observed that when MANT-ATP was preincubated with PCNA, the addition of RFC led to a larger increase in FRET signal within the dead time (1.31x over baseline, like before), followed by a minor increase in fluorescence intensity over the course of one hour (1.31 to 1.34x). Adding p/t-DNA to this sample rapidly decreased fluorescence during the dead time of mixing (from 1.34x to 1.30x), followed by a decay in signal at the same rate as in the first time course (from 1.30x to 1.24x). This result further suggests that the decay is due to MANT-ATP hydrolysis.

Collectively, our results indicate that RFC exhibits slow nucleotide exchange that is converted to rapid exchange upon interaction with PCNA. Our results also indicate that interaction with p/t-DNA can somewhat accelerate exchange, but not to a biologically relevant timescale necessary for eficient clamp loading. Thus, PCNA is a nucleotide exchange factor for RFC.

#### MD simulations predict that PCNA promotes nucleotide exchange in RFC by opening the D-E interface

To investigate how PCNA might promote nucleotide exchange in RFC, we ran additional all-atom molecular dynamics simulations. Based on our cryo-EM reconstructions and biochemical data, we performed five 2-μs long simulations of an autoinhibited RFC:PCNA complex where the A, B, and C subunits are ATP-bound and the D subunit is ADP-bound (the E subunit remained GDP-bound).

In the autoinhibited state, the A, B, and C subunits contact PCNA while the D and E subunits do not (12, 24, 28). The D and E subunits need to bind to PCNA for opening of PCNA (12). In one of our simulations, the D subunit tightly engages PCNA before the E subunit does (**Fig. 6A & Fig. 6S1**). This interaction pries the D subunit away from the E subunit because the tight AAA+ spiral of autoinhibited RFC is geometrically incompatible with full PCNA engagement. The opening of the D-E interface exposes the D subunit’s active site, leading to spontaneous dissociation of ADP in the simulation (**Fig. 6A**). Rotation of the D subunit’s lid subdomain upon ADP release sampled in our simulations is similar to the rotation we observe in the C subunit upon ADP release in our cryo-EM reconstructions (**Fig. 3 & Fig. 6B,C**). Thus, across all our reconstructions and simulations, C and D exhibit similar nucleotide-dependent conformational changes, which may explain why these two subunits exchange ADP slowly. ADP release is sampled only in this trajectory among our replicates: in three of our simulations, D and E make little-to-no contacts with PCNA, and in the remaining simulation, the D and E subunits concomitantly engage PCNA (**Fig. 6S1**).

**Figure 6.**
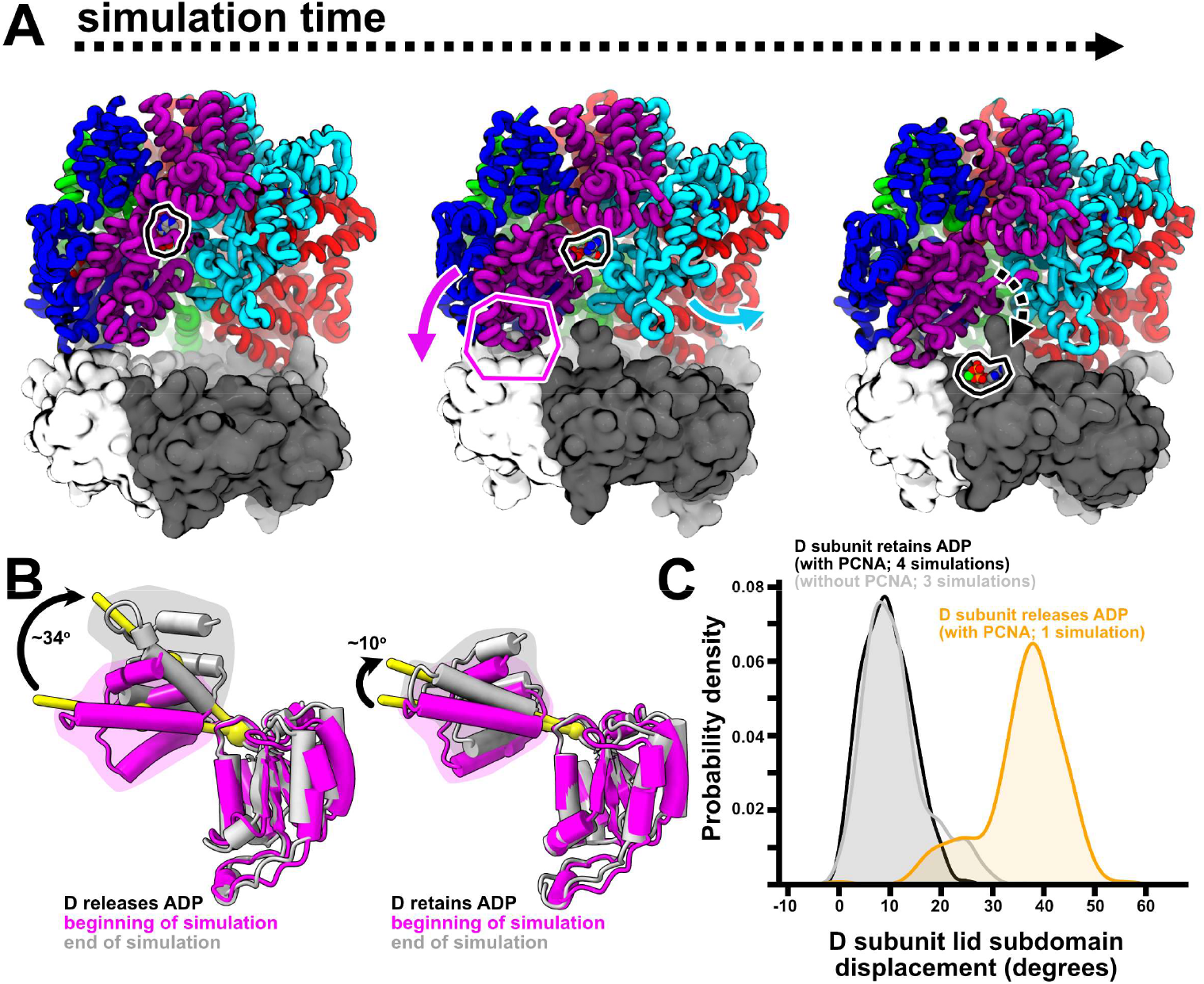
MD simulations predict how PCNA promotes ADP release. **A**. Snapshots from an equilibrium simulation show that as the ADP-bound D subunit engages with PCNA, it is pried away from the E subunit and ADP is released from its active site. **B**. ADP release from the D subunit coincides with a large rotation of the lid subdomain in the same direction observed in the experimentally determined structure of the C subunit (compare with **Fig. 3A**). **C**. Sampled distribution of the D subunit’s lid subdomain rotation from equilibrium simulations of ADP-bound RFC without PCNA (silver), RFC:PCNA where ADP is not released (black), and RFC:PCNA where ADP is released (orange) is shown as kernel density estimates.

## Discussion

RFC uses energy from the ATPase cycle to load PCNA onto DNA during replication and damage repair. RFC can load PCNA onto p/t-DNA with a maximal velocity of 1 s^-1^ (6, 10). Here, our cryo-EM reconstructions and biochemical data show that, on its own, RFC counter-intuitively releases ADP much slower than these measured rates, on the scale of hours to days. The apparent discrepancy between measured rates of clamp loading and our structural and biochemical data is resolved by PCNA, which promotes ADP release from RFC on the timescale required for clamp loading.

### PCNA is a nucleotide exchange factor for RFC

To load PCNA at the necessary rate to support DNA replication and repair, ADP release from RFC needs to be catalyzed such that it is not rate-limiting. Our biochemical and molecular dynamics data indicate that PCNA itself acts as a nucleotide exchange factor for RFC by promoting ADP release from the D subunit. As the D subunit engages with PCNA, the D-E interface is weakly bridged by ADP, making it susceptible to opening for ADP release. While we did not directly address whether PCNA helps promote exchange at the C subunit, we speculate that an analogous mechanism could pry apart the ADP-bound C-D interface as C binds PCNA.

Our data agrees with previous findings that RFC binds one or two additional ATP molecules after PCNA binding (10, 33). However, these previous measurements could not explain how RFC was precluding ATP binding until it bound PCNA, pinpoint which RFC subunits were binding the additional ATP molecules, nor explain how PCNA promoted additional ATP binding. Our data addresses all three of these questions by showing that RFC precludes ATP binding by releasing ADP slowly from the C and D subunits, and that PCNA promotes additional ATP binding by catalyzing ADP release from RFC.

### PCNA’s role as a nucleotide exchange factor may order the clamp loading reaction

RFC’s interaction with PCNA is ATP-dependent (10, 35). We speculate that fast ADP release and ATP binding to the A and B subunits (and possibly C) stabilizes their AAA+ domains in a conformation competent for PCNA binding (**Fig. 7A**). PCNA binding then promotes ADP release in the D subunit (and possibly C). PCNA-induced ATP binding completes the binding interface between AAA+ subunits, driving the opening of PCNA and subsequent DNA binding. If these interfaces were completed prior to binding PCNA, RFC may aberrantly expose its DNA binding chamber and bind DNA before PCNA, which is a dead-end, off-pathway process. We propose that by using its own substrate as a nucleotide exchange factor, RFC can limit off-pathway configurations that may arise from binding DNA prior to PCNA.

**Figure 7.**
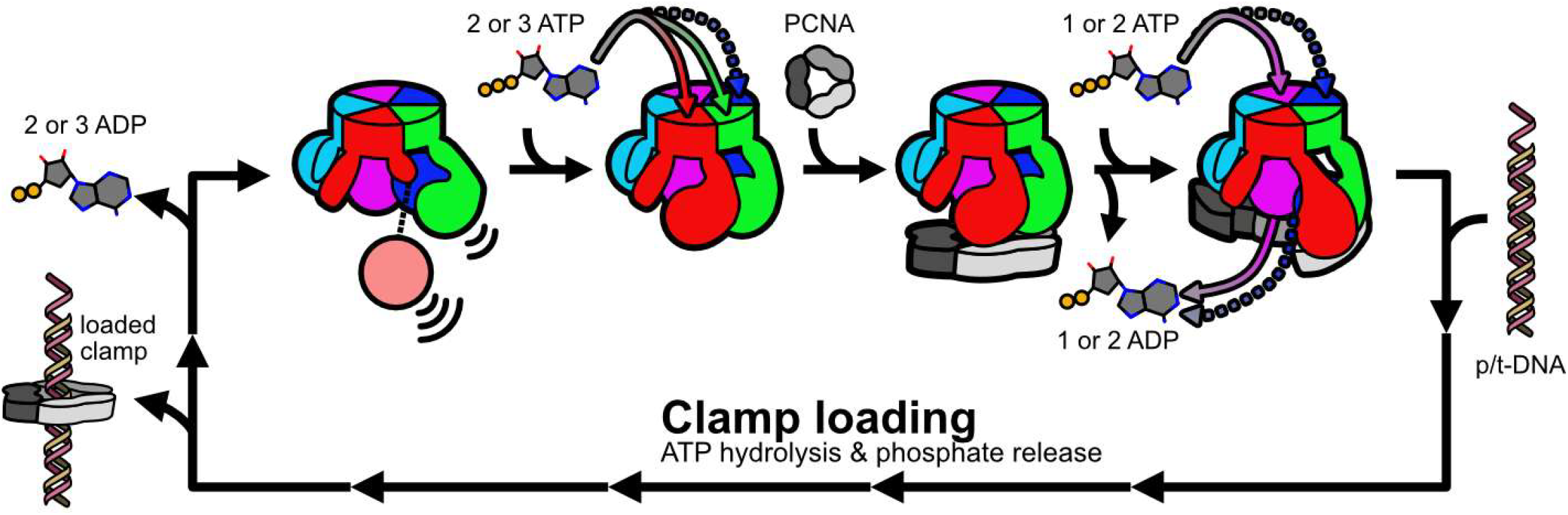
A speculative model for the role of nucleotide exchange in RFC. Mixed nucleotide occupancy RFC has flexible apo subunits (A and B) and more rigid ADP-bound subunits (C and D). Two or three ATP molecules binding at A, B, and possibly C stabilizes RFC in a conformation competent to bind PCNA. PCNA binding promotes additional ADP release from the remaining subunits, and ATP binding drives crab-claw opening and downstream clamp loading.

### Connection to other clamp loaders

Eukaryotes have at least three other ‘alternative’ clamp loaders (36). Ctf18-RFC loads PCNA onto the leading strand (13, 37), Elg1-RFC (ATAD5-RFC in human) unloads PCNA (38), and Rad24-RFC (Rad17-RFC in humans) loads the alternative sliding clamp 9-1-1 (21, 22). These alternative loaders are formed by replacing the A subunit of RFC with the eponymous protein, while the B, C, D and E subunits remain unchanged. Hence, it is possible that PCNA can promote nucleotide exchange in the D subunit of these alternative loaders as well. Recently determined structures of ATAD5-RFC bound to PCNA found that the D subunit can be ATP-γS-bound or ADP-bound depending on the conformational state (20). While the ordering of these intermediate states is ambiguous (*i*.*e*., one cannot distinguish if ATP-γS hydrolysis led to the ADP-bound intermediate or if nucleotide exchange led from the ADP-bound to the ATP-γS-bound intermediate), these structures again highlight the unique plasticity of the RFC D subunit. These results suggest that the loader’s interaction with PCNA is somehow connected to its nucleotide occupancy.

Like the eukaryotic loaders, bacterial and phage clamp loaders are also hetero-pentameric AAA+ assemblies, and not every subunit is necessarily an active ATPase. It was previously proposed that the *E. coli* clamp loader binds two ATP molecules in absence of its sliding clamp, with the final ATP molecule binding upon clamp binding (39). Our recent structural work showed that the *E. coli* clamp loader opens its sliding clamp primarily through conformational changes at its C subunit (40). Thus, we propose that if the *E. coli* sliding clamp also promotes nucleotide exchange in its clamp loader, it most likely occurs at the C subunit.

### Nucleotide exchange in other ATPases

Although RFC shares many similarities with the broader class of the AAA+ family of ATPases, it is also somewhat unusual. Most AAA+ ATPases are homo-hexameric ring-shaped assemblies that use energy from ATP to translocate a substrate through their central pore (41–43). These assemblies typically use a hand-over-hand mechanism, creating a cracked interface or ‘seam’ which exposes one subunit’s ATPase active site. ADP release and ATP binding are thought to readily occur at the exposed site and so it is unlikely that these systems require additional factors to promote nucleotide exchange. ATP hydrolysis and nucleotide exchange proceeds around the AAA+ ring in a rotary fashion, giving rise to a continuous translocation mechanism. In contrast, RFC acts as a ‘one-and-done’ protein remodeling switch that opens and closes PCNA around DNA. Because RFC loads PCNA through a crab-claw mechanism and not a hand-over-hand mechanism (12), it does not create a seam between adjacent ATPase sites. ADP release is promoted by either flexibility of subunits when RFC is not engaged with its substrates (*e*.*g*., the A and B subunits), or from an exchange factor which pries apart subunit interfaces (*e*.*g*., PCNA and the D/E interface).

The need for external factors that promote nucleotide exchange may be common amongst ATPases that act as switches. Hsp70 and DnaA are two such ATPases that toggle on or off based on their nucleotide-bound state. Hsp70 is a chaperone which binds unfolded protein substrate (44, 45), and DnaA is a switch that regulates DNA replication initiation (46, 47). In both cases, conversion from the ADP-bound state back to the ATP-bound state requires interactions with accessory factors. In *E. coli*, specific DNA sequences called DnaA Reactivating Sequences bind ADP-bound DnaA and promote ADP release (48). For Hsp70, nucleotide exchange is catalyzed by the binding partner GrpE, which wedges Hsp70’s nucleotide binding cleft open to promote ADP release (49, 50).

### Analogy to GTPases, GEFs, and GAPs

The relationship between RFC and its substrates is reminiscent of how small GTPases interact with regulatory proteins Guanine Nucleotide Exchange Factors (GEFs) and GTPase-Activating Proteins (GAPs). Like RFC, most small GTPases are molecular switches that are in the ‘on’ state when nucleotide-triphosphate bound and in the ‘off’ state when nucleotide-diphosphate bound (51). In these systems, GEF binding promotes GDP release, and GAP binding catalyzes GTP hydrolysis (52, 53). In this way, PCNA can be thought of as a nucleotide exchange factor (NEF) analog to GEF, and DNA can be thought of as the NTPase activating partner (NAP) analog to GAP.

A stark difference between RFC and other switch-like NTPases is that these other systems typically act on one substrate at a time. Not only does RFC need to act on two substrates simultaneously, but it also has the added constraint that it needs to bind PCNA before it binds DNA. At face value, these constraints increase the complexity of RFC’s activity. However, they also afford an opportunity for RFC to simplify the choreography of its ATPase cycle. The other switch-like NTPases typically use dedicated NEFs and NAPs that are independent of their substrates. For instance, Hsp70’s substrate is unfolded protein, but it uses GrpE as its NEF and DnaJ as its NAP (45). On the other hand, RFC’s two substrates moonlight as its NEF and NAP, reducing the overall number of components and complexity of the system and potentially providing a defined order to the clamp loading reaction.

## Methods

### Protein expression and purification

#### RFC

RFC was purified as described previously (12, 54), with some modifications. pET(11a)-RFC[2 + 3 + 4] and pLANT-2/RIL-RFC[1 + 5] were co-transformed into either BL21(DE3) or Rosetta(DE3) *E. coli* cells (Millipore). Transformants were grown in preculture overnight and used to inoculate 6 L of terrific broth medium supplemented with 50 μg/mL kanamycin and 100 μg/mL ampicillin at 37 oC and shaking at 200 rpm. Once the cultures reached an optical density of ∼1, they were cold-shocked for twenty minutes by incubating at 4 °C, and then protein expression was induced with IPTG. These cultures were then shaken at 200 rpm at 18 oC for ∼16 hours to express protein. Cultures were pelleted and pushed through a syringe directly into liquid nitrogen to flash freeze as noodles. Noodles were cryo-milled with a planetary ball mill grinder at liquid nitrogen temperature. Cryo-lysed cells were resuspended with extraction buffer containing 30 mM [4-(2-hydroxyethyl)piperazin-1-yl]ethanesulfonic acid (HEPES)–NaOH (pH 7.5), 250 mM NaCl, 0.25 mM Ethylenediaminetetraacetic acid (EDTA), 5% (v/v) glycerol, 2 mM Dithiothreitol (DTT), 2 μg/ml aprotinin, 0.2 μg/ml pepstatin, 2 μg/ml leupeptin, and 1 mM 4-(2-aminoethyl)benzenesulfonyl fluoride hydrochloride (AEBSF).

RFC was purified by chromatography over a 10 mL HiTrap SP Sepharose FF column (Cytiva) using an 80 mL gradient from 300 to 600 mM NaCl. Peak fractions were pooled and purified by size exclusion chromatography on a HiLoad 16/600 Supradex 200 pg (Cytiva) in storage buffer containing 25 mM Hepes-NaCl (pH 7.5), 300 mM NaCl, 2 mM DTT, and 5% (v/v) glycerol. RFC was concentrated using a 100 kDa MWCO Amicon Ultra-15 Centrifugal Filter Unit (EMD Millipore), flash frozen in liquid nitrogen, and stored at -80 oC.

#### PCNA

PCNA was purified as described previously (52). pET-28 vector encoding yeast PCNA with a Prescission protease cleavable N-terminal 6xHis tag was transformed into BL21(DE3) *E. coli* cells. Transformants were grown in preculture overnight and used to inoculate 1 L of terrific broth medium supplemented with 50 μg/mL kanamycin. The culture was grown at 37 oC and 200 rpm to an optical density at 600 nm of ∼1. The culture was then cold-shocked by incubation at 4 °C for twenty minutes. Protein expression was induced with addition of IPTG to a final concentration of 1 mM, and grown overnight at 18 oC. Cells were pelleted and resuspended in lysis buffer containing 30 mM HEPES-NaOH (pH 7.6), 20 mM imidazole, 500 mM NaCl, 10% (v/v) glycerol, and 5 mM β-mercaptoethanol.

The resuspended cells were lysed with a cell homogenizer (Microfluidics Inc), centrifuged, and filtered through a 0.45 μm filter before applying to a 5 mL HisTrap FF column (Cytiva). The column was washed with a high salt buffer (lysis buffer with 1 M NaCl) and then a low salt buffer (50 mM NaCl). PCNA was eluted with a 50% step of 500 mM imidazole. The His tag was cleaved with Precision protease for two hours at room temperature. Cleaved protein was applied to a 5 mL HiTrap Q HP column (Cytiva) and eluted with a 2 M NaCl buffer 100 mL gradient. Peak fractions were collected and dialyzed against buffer containing 30 mM Tris (pH 7.5), 100 mM NaCl, and 2 mM DTT. PCNA was concentrated using a 30 kDa MWCO Amicon Ultra-15 Centrifugal Filter Unit (EMD Millipore), flash frozen in liquid nitrogen, and stored at -80 oC.

#### E. coli Beta clamp

β-clamp expression plasmid is as described in (53) and the protein was purified as described in (37). β-clamp with a Prescission protease cleavable N-terminal 6-His tag was expressed in BLR21(DE3) *E. coli* (Millipore). Transformants were precultured overnight and used to inoculate 1 L of terrific broth media supplemented with 50 μg/mL kanamycin and grown at 37 oC until the culture reached an optical density between 0.6 to 0.8. Then the culture was cold shocked by incubation at 4 °C for twenty minutes. Protein expression was induced with addition IPTG (1mM final), and the culture was grown overnight at 18 oC. Cultures were pelleted, flash frozen in liquid nitrogen, and stored at -80 oC until purification.

The frozen pellet was thawed on ice and resuspended in extraction buffer containing 20 mM Tris (pH 8.0), 500 mM NaCl, 10% (v/v) glycerol, 5 mM β-mercaptoethanol, and 20 mM imidazole, and lysed with a cell homogenizer (Microfluidics Inc). The lysate was clarified by centrifugation and loaded onto HisTrap columns (Cytiva) and washed with five column volumes of extraction buffer. The protein was eluted with buffer containing 20 mM Tris (pH 8.0), 500 mM NaCl, 10% (v/v) glycerol, 5 mM β-mercaptoethanol, and 250 mM imidazole. Peak fractions were pooled and dialyzed against buffer containing 50 mM Tris (pH 8.0), 10% (v/v) glycerol, and 2 mM DTT while also treated with Precision protease to cleave the N-terminal His tag. The protein was loaded onto a 5-mL HiTrap Q column (Cytiva) and eluted with a gradient reaching a final salt concentration of 500 mM NaCl. Peak fractions were pooled and dialyzed against buffer containing 50 mM Tris (pH 7.5), 50 mM NaCl, 10% (v/v) glycerol, and 2 mM DTT. Lastly, purified β-clamp was concentrated with a 30 kDa MWCO Amicon Ultra-15 Centrifugal Filter Unit (EMD Millipore), flash frozen in liquid nitrogen, and stored at -80 oC.

### Cryo-EM sample preparation

Purified RFC was buffer exchanged from its storage buffer into buffer containing 25 mM Hepes-KOH (pH 7.5), 300 mM K(OAc), 7 mM Mg(OAc)_2_, 1 mM TCEP, and 5% (v/v) glycerol, and diluted to 2 mg/mL. UltrAuFoil R 2/2 grids were glow discharged in a Pelco easiGlow for 60 s at 25 mA (negative polarity). 3.5 μL of sample was applied in a Vitrobot Mark IV (FEI) maintaining 4 oC and 100% humidity. Grids were blotted after a 5 s wait time, with a blot force of 2 and blot time of 30 s. The grids were plunged into liquid ethane and stored in liquid nitrogen until collection.

### Cryo-EM data collection

RFC was imaged on a Titan Krios operating at 300 kV. The Krios was equipped with a GIF energy filter, and a pixel size of 0.415 Å using a K3 Summit detector in super-resolution counting mode. The data was collected in one session with a total exposure dose of 49.9 e^-^/Å^2^ per micrograph averaging 34 frames. The resulting collection yielded a total of 7,660 micrographs.

### Cryo-EM data processing and reconstruction

All processing was done using CryoSPARC version 4.X (54) (**Fig. 1S1-1S4**). Micrographs were initially aligned with Patch Motion Correction (F-crop factor: ½) and CTF estimation using Patch CTF. The micrographs were denoised using CryoSPARC’s Micrograph Denoiser job; these are used for visualization of the micrographs and particle picking, but particles were extracted from the unprocessed micrographs. Blob picking (diameter range 70 to 140 Å) was used on a subset of 500 micrographs. These particles were extracted in a box size of 256 pixels and Fourier cropped to 128 pixels. Initial 2D classification of these particles into 100 classes yielded high quality class averages with visible secondary structural elements. These 2D classes were then used as templates to pick on all 7,660 micrographs (diameter 90 Å). We picked a total of 7.1 million particles for downstream classification. Particles were 2D classified into 250 classes. 48 selected classes were further classified into 100 2D classes. 36 selected classes were chosen for downstream processing, approximately 2.4 million particles.

After ab initio refinement, we performed iterative 3D classification into six final 3D classes. Each class was subjected to non-uniform refinement (55). Maps were post-processed with EMReady (56). Visualizations in this manuscript depict maps after processing with EMReady, and a comparison between the unprocessed and processed maps can be found in **Fig. 1S5**.

### Model building and refinement

Starting models of RFC taken from the autoinhibited RFC:PCNA complex (PDB: 7thj) were rigid-body fit into the unprocessed cryo-EM maps. The models were initially refined in ISOLDE (55) using both the EMReady processed maps and the unprocessed maps to guide the eye. After, the models were manually adjusted in Coot (56, 57) using both the EMReady processed and unprocessed maps, and finally refined in Phenix (58) using only the unprocessed maps. UCSF ChimeraX was used to visualize maps and models (59, 60).

### Equilibrium molecular dynamics simulations

Starting structures of RFC were taken autoinhibited RFC:PCNA complex (PDB: 7thj). To simulate RFC alone, the three chains of PCNA were deleted; ATPγS was manually changed to either ATP or to ADP by deleting the γ-phosphate. Structures were then prepared for simulation with the CHARMM-GUI webserver (61). The CHARMM36m force field (62) with CUFIX nonbonded corrections (63) was used to describe protein interactions. The protein complex was solvated in an octahedral box with 14 Å padding, and water was described with the CHARMM-modified TIP3P model (64). Hydrogen mass repartitioning was applied to allow for a 4-fs timestep to propagate the equations of motion (65). Simulations were performed with the GPU-accelerated NAMD3 engine (66). All necessary topology files, starting coordinate files, and NAMD input files are included in the supplemental information.

The systems were subjected to 10,000 steps of conjugate gradient energy minimization, followed by slow equilibration in three steps. First, 125,000 2-fs timesteps (250 ps total) were performed in the NVT statistical ensemble while restraining all heavy atoms (1.0 scaling for backbone; 0.5 scaling for side chains). Bonds involving hydrogen were constrained with the SHAKE algorithm (67). Temperature was reassigned every 500 timesteps to the target 310.15 K. Next, 125,000 2-fs timesteps (250 ps total) were performed in the NPT statistical ensemble, maintaining the pressure with a Langevin Piston barostat (oscillation period 50 fs; oscillation decay 25 fs) and the temperature with Langevin Damping (damping coeficient 1 ps^-1^), constraining heavy atoms as before. Lastly, 125,000 4-fs timesteps (500 ps total) were performed in the NPT statistical ensemble as before, with heavy atom constraints scaled by 0.5. Production simulations were performed in the NPT statistical ensemble with a 4-fs integration timestep and no constraints on the protein for 2 microseconds each. Long range non-bonded interactions were corrected with the Particle-Mesh-Ewald (PME) scheme (68).

### τ -Random Acceleration Molecular Dynamics simulations

τ-Random Acceleration Molecular Dynamics (τ-RAMD) were performed as implemented in NAMD (32). τ-RAMD simulations probe ligand dissociation rates by applying a constant force in a random direction to a ligand. If the ligand’s center-of-mass does not move farther than a threshold distance relative to the center-of-mass of the protein to which it is bound in a set amount of time, a new direction is chosen to apply to apply the force.

Starting structures were seeded from the final frame of each of the independent equilibrium simulations described above. This allowed for sampling exit pathways of ADP from different AAA+ domain arrangements. From each starting structure, ten τ-RAMD simulations were performed for each bound ADP, for a total of 120 independent simulations. A randomly directed force of 14 kcal/mol/Å was applied to ADP, and a threshold distance of 0.025 Å was checked every 50 simulation steps. Simulations were stopped either when the ADP was more than 40 Å away from the center-of-mass of its subunit’s Rossmann fold, or after one day of wall-time, whichever occurred first.

### Simulation visualization and analysis

Simulations were visualized with VMD 1.94a (69). Centers-of-mass distances were calculated with the *colvars* module in VMD (70). The number of contacts between subunits/PCNA were calculated with the MDAnalysis package (71, 72).

### MANT-ATP fluorescence assays

RFC and PCNA were buffer exchanged from their storage buffer into buffer containing 25 mM Hepes-KOH (pH 7.5), 200 mM K(OAc), 7 mM Mg(OAc)_2_, 1 mM TCEP, and 5% (v/v) glycerol. MANT-ATP was purchased from Jena Bioscience. Because we observed mixed nucleotide occupancy in our cryo-EM reconstructions, we incubated concentrated RFC (∼6.5 μM) with 240 μM ADP prior to adding to the assay. 0.3 μM RFC was added to 12 μM MANT-ATP, which was either by itself or incubated with 1 μM PCNA, 1 μM p/t-DNA, or both. MANT-ATP binding to RFC was monitored with Förster Resonance Energy Transfer (FRET) between tryptophan residues of RFC as the donor (λ_EX_ = 297 nm) and MANT-ATP as the acceptor (λ_EM_ = 440 nm). Intensity was read after mixing on a Jobin-Yvon Horiba Fluoromax with slit widths 2 nm (excitation) and 3 nm (emission) with a 1 second integration time. For the 16-hour time courses, timepoints were taken every 6 minutes; for the shorter time courses, timepoints were taken every minute. The temperature was maintained at 25 oC. The average signal intensity from the five minutes leading up to addition of RFC was used to determine the baseline signal intensity.

## Supporting information

molecular dynamics input files

RFC models

## Data availability

Cryo-EM maps and models are deposited to the EMDB and PDB, respectively, and will be released upon peer-review and final publication. In the interim, refined models are available as supplemental files and unprocessed/EMReady processed maps are available upon request. All input files required for molecular dynamics simulations are included in the supplement.

## Acknowledgements

We thank Dr. David Lambright for guidance on setting up the MANT-ATP FRET assay. We thank Dr. Bill Royer for use of his size exclusion column during protein purification. We thank Dr. Natasha Buwa for guidance on cryo-milling. We thank Drs. Ala Shaqra and Jeong Min Lee for guidance on size exclusion chromatography. We thank Drs. KangKang Song, Christna Ouch, and Jeng-Yih Chang at the UMass Chan cryo-EM facility for help collecting the cryo-EM data. We thank Dr. Emma Sedivy and other members of the Kelch lab for helpful discussions during preparation of this manuscript.

This work was funded by American Cancer Society award PF-22-114 to J.P. and NIGMS awards R35GM156361, R01-GM127776, and R01-GM145943 B.A.K.. Cryo-EM data was collected at the UMass Chan cryo-EM facility, which was supported by a grant from the Massachusetts Life Science Center. This research used the Delta advanced computing and data resource which is supported by the National Science Foundation (award OAC 2005572) and the State of Illinois. Delta is a joint effort of the University of Illinois Urbana-Champaign and its National Center for Supercomputing Applications. The allocation for Delta was awarded through the NSF ACCESS mechanism, which is supported by National Science Foundation (Grants ACI-2138259, 2138286, 2138307, 2137603, and 2138296).

Computational resources were also provided by UMass Chan Scientific Computing for Innovation (SCI) cluster.

## Author contributions

J.P. and B.A.K. conceptualized research and analyzed data. J.P. purified RFC, performed all cryo-EM sample preparation, cryo-EM data processing, model building, molecular dynamics simulations, and MANT-ATP FRET assays. J.L. purified β-clamp and assisted with conceptualization. X.L. purified PCNA and assisted with conceptualization. K.A. purified PCNA. S.L. assisted with purifying RFC. J.P. and B.A.K. wrote the manuscript.

