## supplemental figures for "PCNA is a Nucleotide Exchange Factor for the Clamp Loader ATPase Complex"

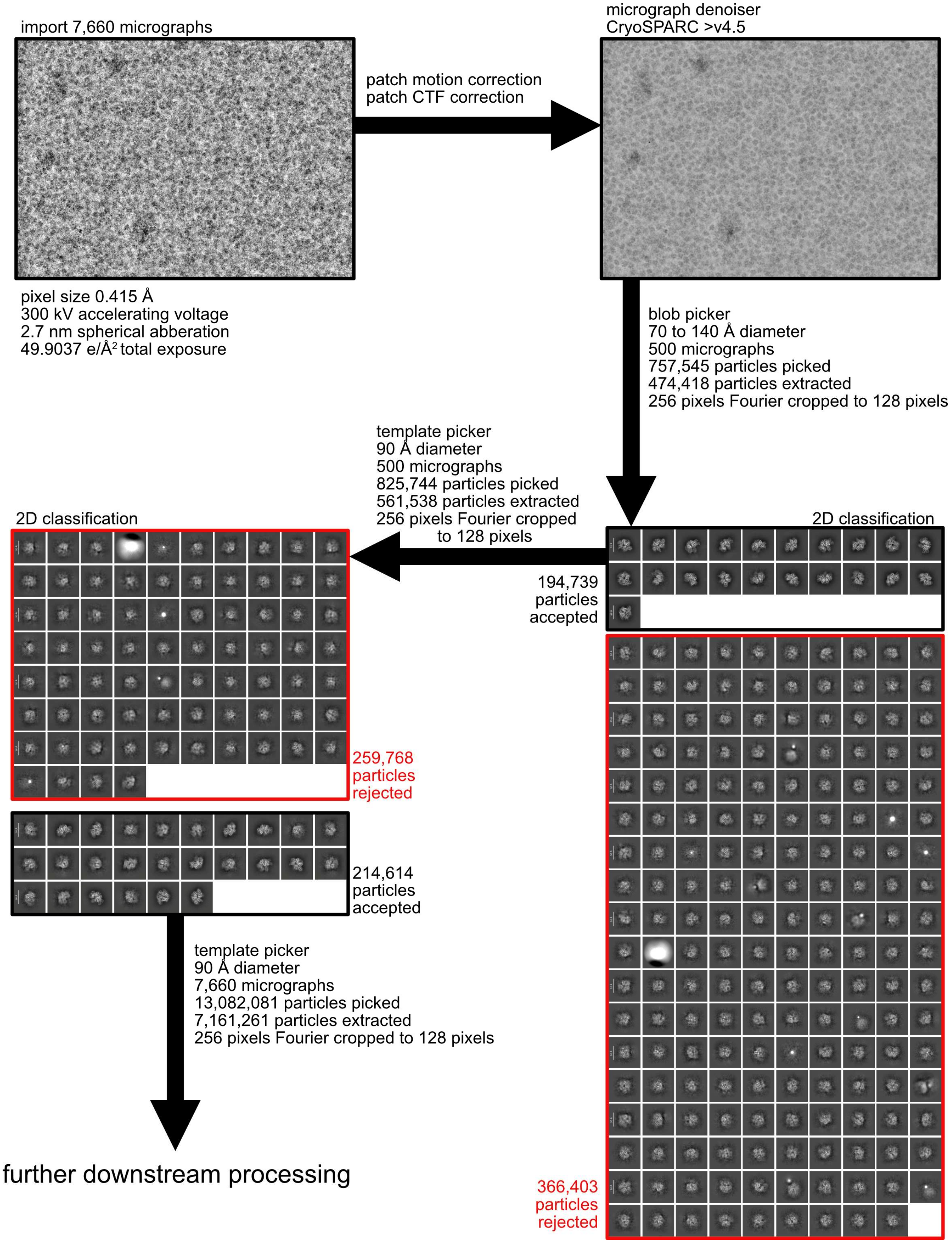


**Supplemental Figure 1S1.** Micrograph processing, particle picking, and initial 2D classification.

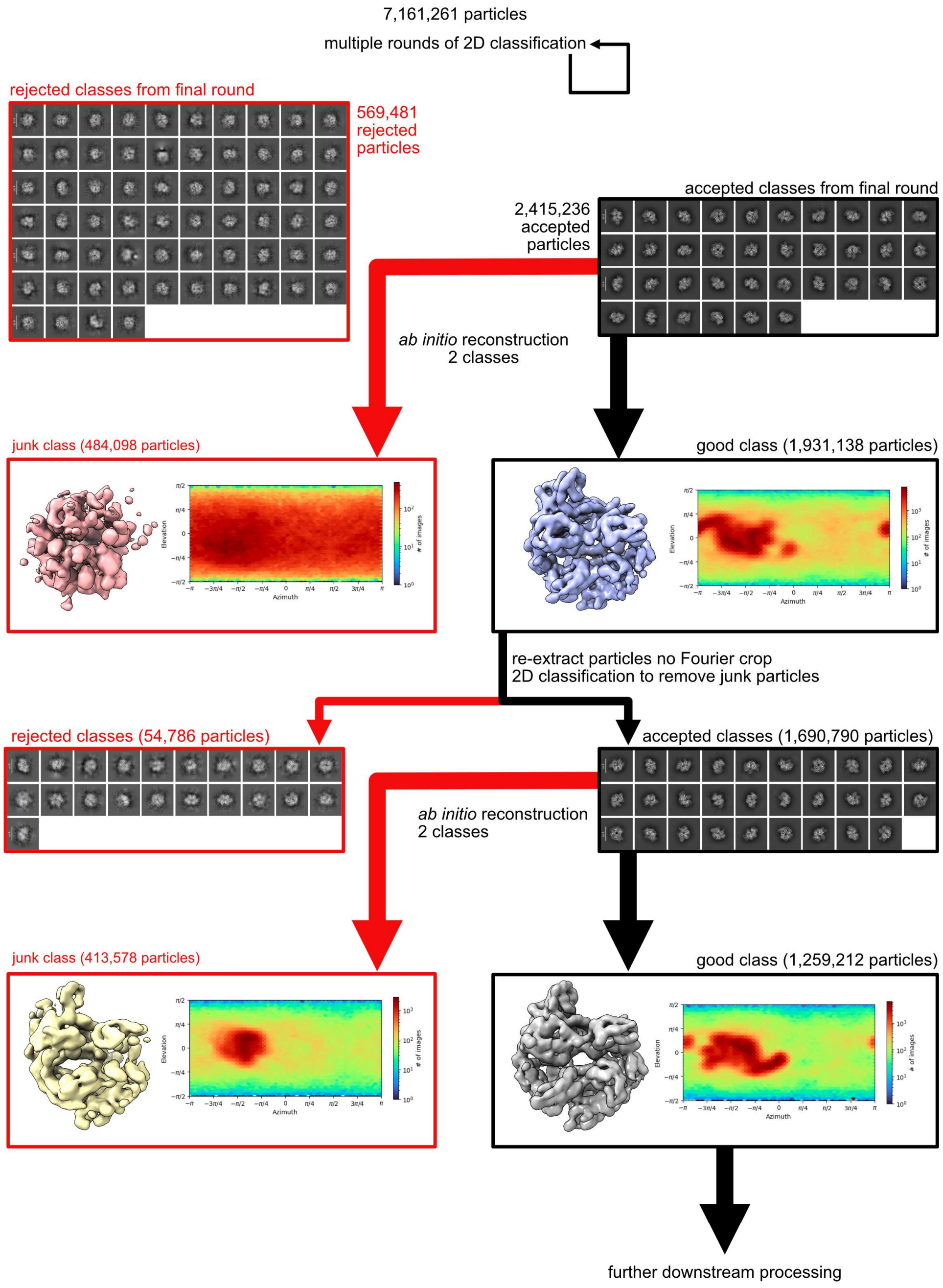


**Supplemental Figure 1S2.** 2D classification, particle curation, and *ab initio* reconstructions.

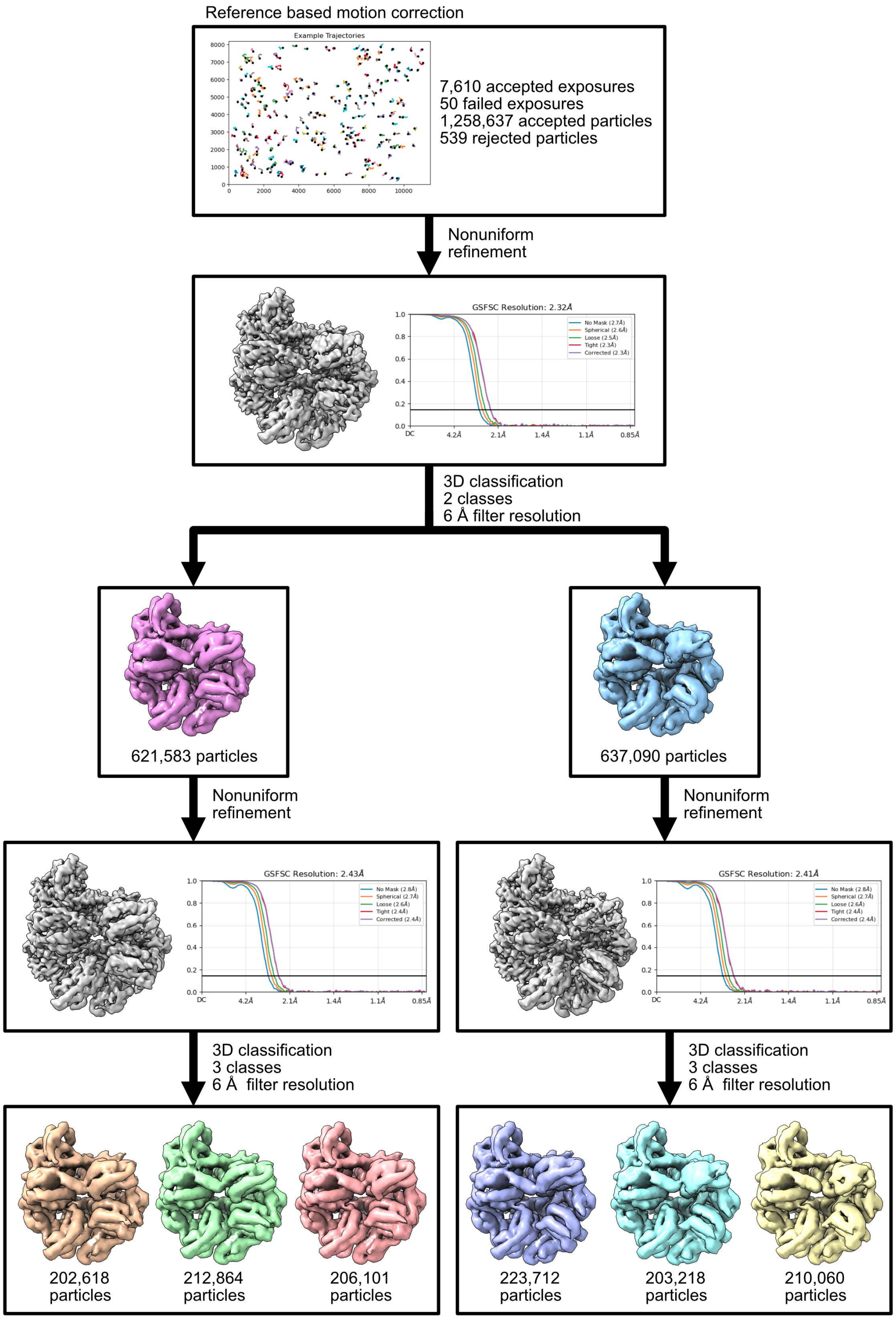


**Supplemental Figure 1S3.** 3D classification and refinement.

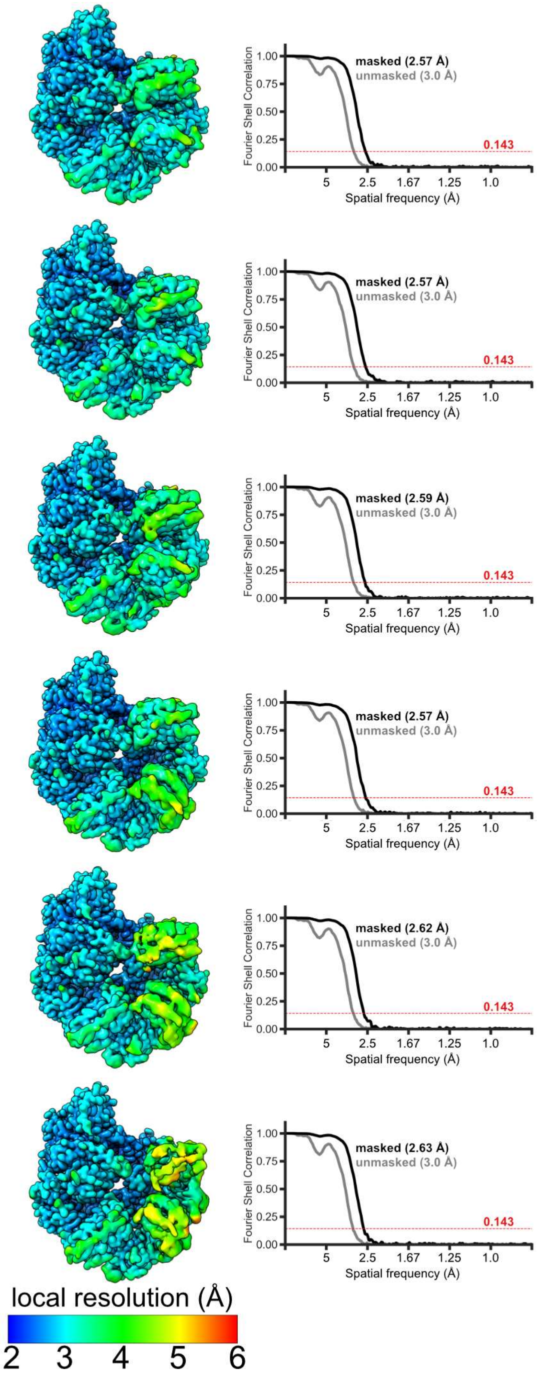


**Supplemental Figure 1S4**. Local resolution and overall resolution by gold-standard FSC for all six final 3D classes.

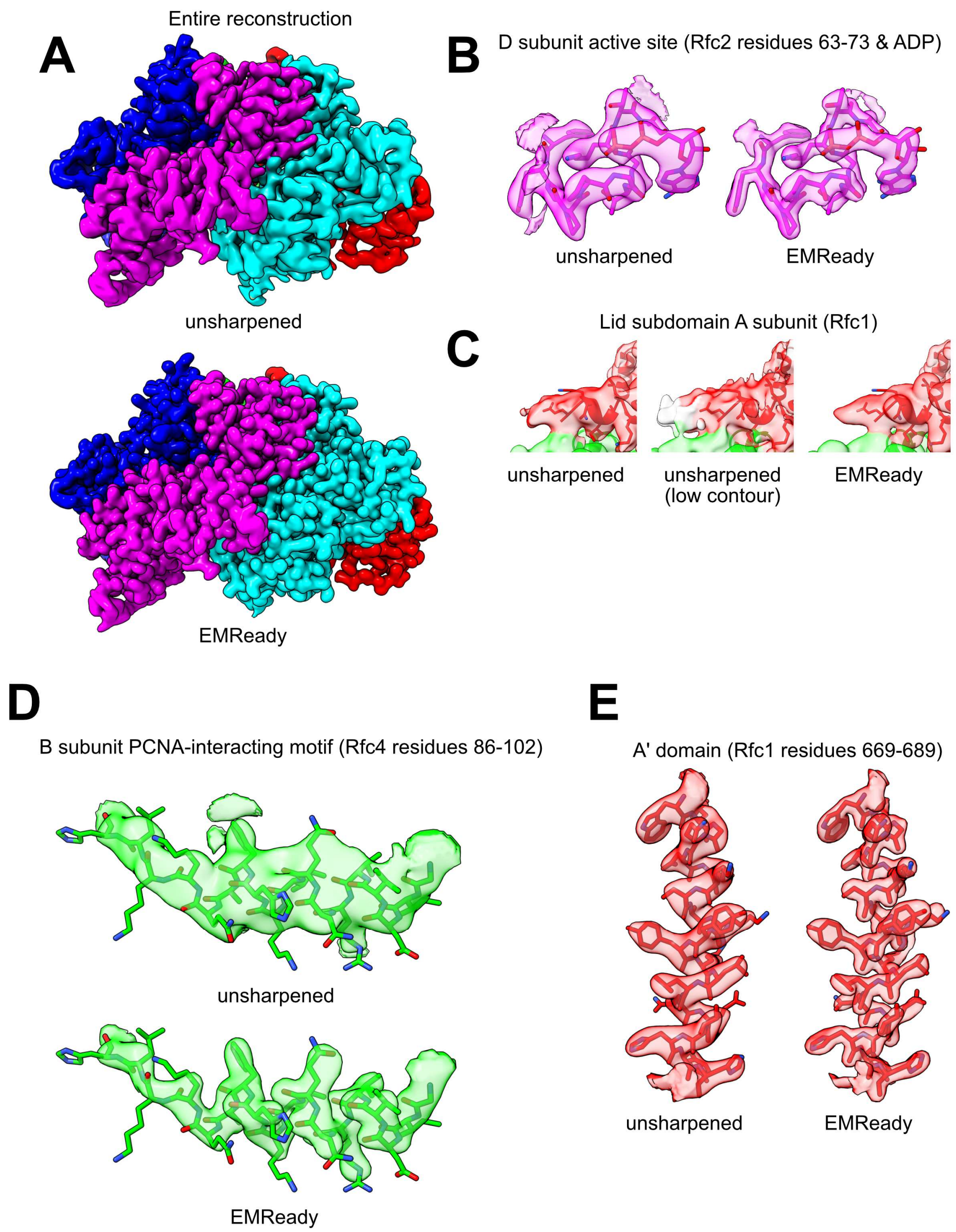


**Supplemental Figure 1S5. Map sharpening and map-to-model fits. A.** One reconstruction shown before and after processing with EMReady. EMReady processed maps were used for figure making and initial model building, but final refinements in Phenix used the unprocessed maps. **B.** ATPase active site of the D subunit before and after processing with EMReady. **C.** Density where the A subunit begins to appear in the unsharpened map at two contour levels and the EMReady processed map. **D.** Portion of the B subunit which contacts PCNA before and after processing with EMReady. **E.** Portion of the A' domain before and after processing with EMReady.

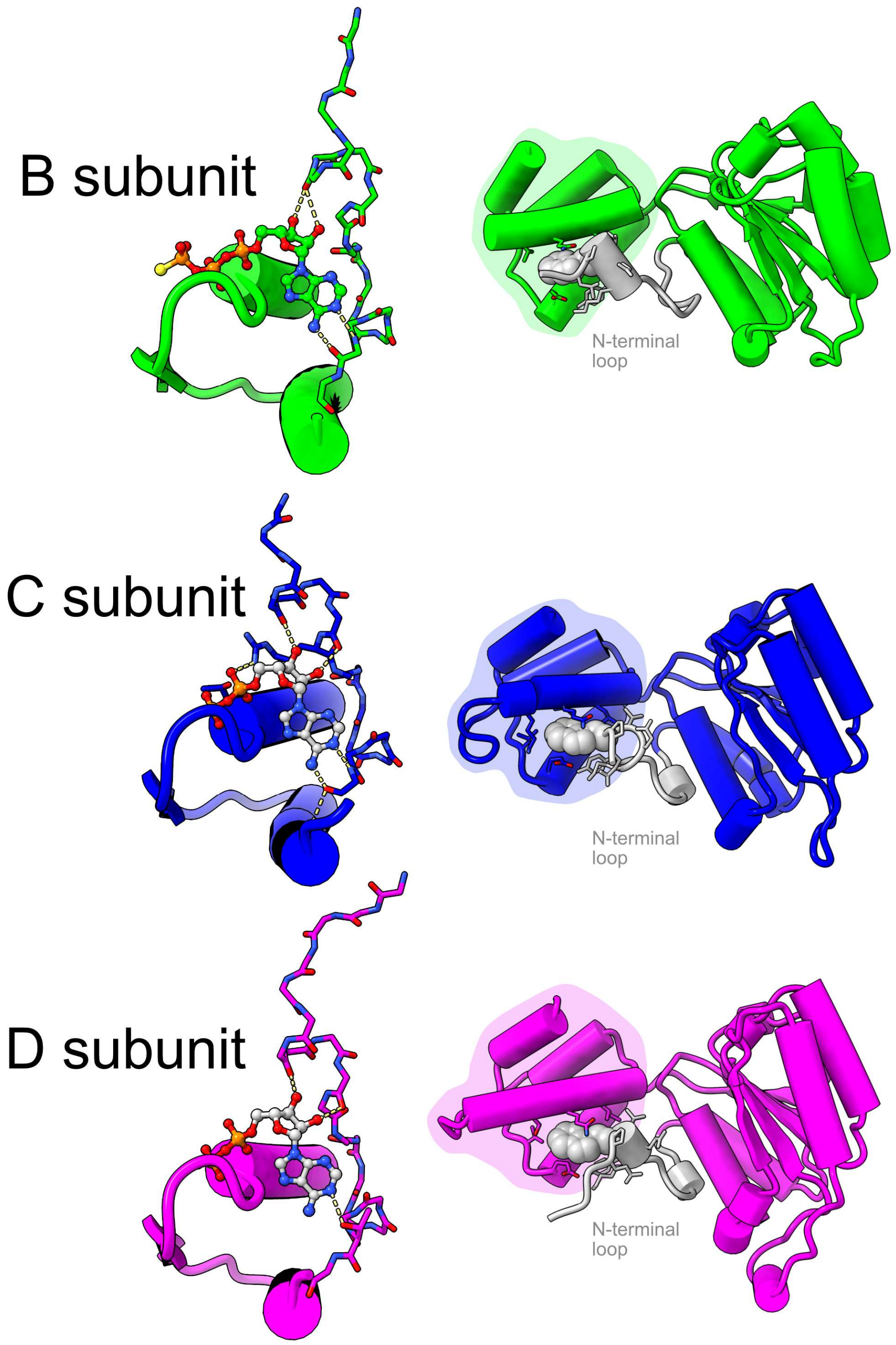


**Supplemental Figure 3S1.** The N-terminal loops of subunits hydrogen bond with nucleotide and interact with the lid subdomain. The autoinhibited state (PDB: 7thj) is used for visualization to show how this interaction occurs in the B subunit, which is apo in all of our reconstructions. (**left**) The modeled N-terminal loop of the B, C, and D subunits is shown as sticks, and hydrogen bonds to nucleotide are shown as dashed yellow lines. Only the backbone of the N-terminal loop is shown for clarity. (**right**) The N-terminal loop is silver, and the lid subdomain is highlighted with a background color. For visualization, a hydrophobic residue of the N-terminal loop is shown as van der Waals spheres, highlighting how it buries into the lid subdomain.

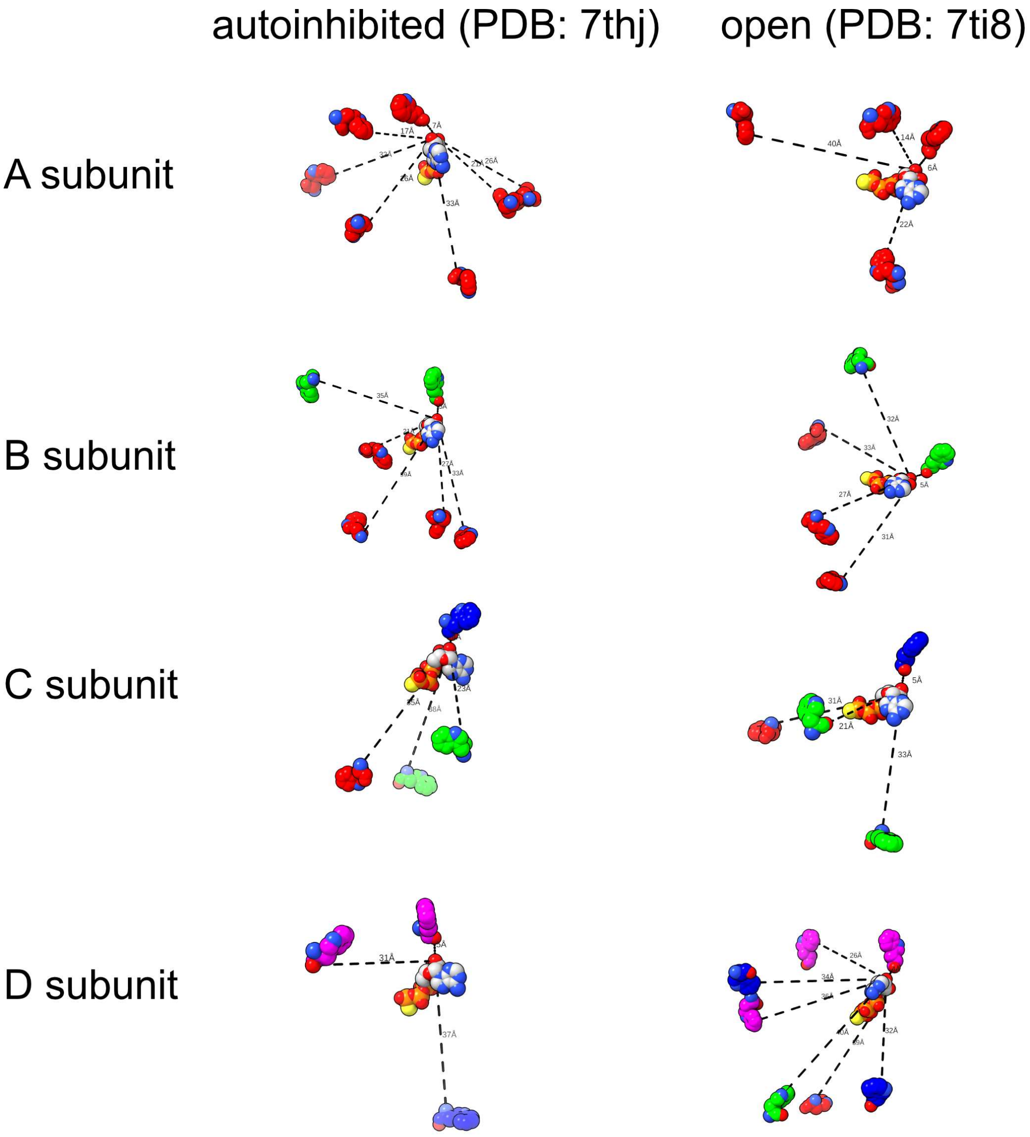


**Supplemental Figure 5S1. Distances between tryptophan residues and ATP.** All tryptophan residues within 30 Å (tryptophan-MANT *R₀* is approximately 25 Å) of bound nucleotide are shown. Distances are reported to the closest oxygen on the ribose sugar, as this is where the MANT moiety attaches. The distances and number of tryptophan residues within the 30 Å cutoff and changes as RFC undergoes the crab-claw conformational change. Nevertheless, there is always at least one tryptophan residue within ~5-7 Å, which will contribute to signal.

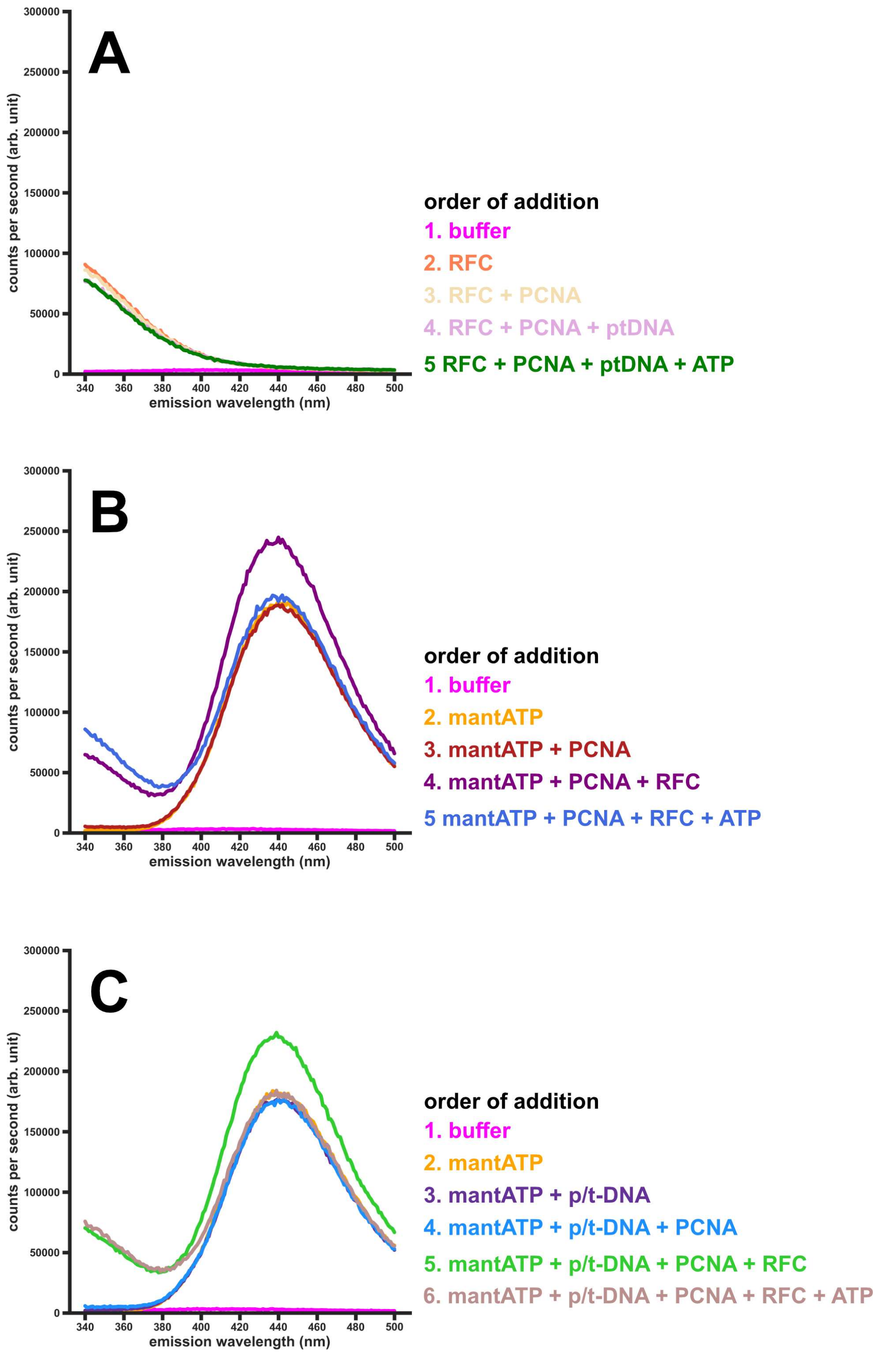


**Supplemental Figure 5S2. Emission spectrum of various conditions. A.** Emission spectra in absence of MANT-ATP. Each spectrum progressively adds a listed component. Signal at 340 nm increases upon the addition of RFC, stays the same upon the addition of PCNA (which does not contain any tryptophan residues), drops slightly upon the addition of p/t-DNA, and stays the same upon the addition of ATP. In all cases, the signal at 440 nm remains largely unaffected **B.** Emission spectra including MANT-ATP and protein components. Signal at 440 nm increases upon the addition of MANT-ATP and stays the same upon the addition of PCNA (which does not bind MANT-ATP). Signal at 440 nm increases ~1.3x upon the addition of RFC, and this signal is quenched upon the addition of excess ATP. **C.** Emission spectra including MANT-ATP, protein components, and DNA. As before, the addition of MANT-ATP increases signal at 440 nm, which stays the same upon the addition of p/t-DNA and PCNA. Signal at 440 nm increases ~1.3x upon the addition of RFC, and this signal is quenched upon the addition of excess ATP.

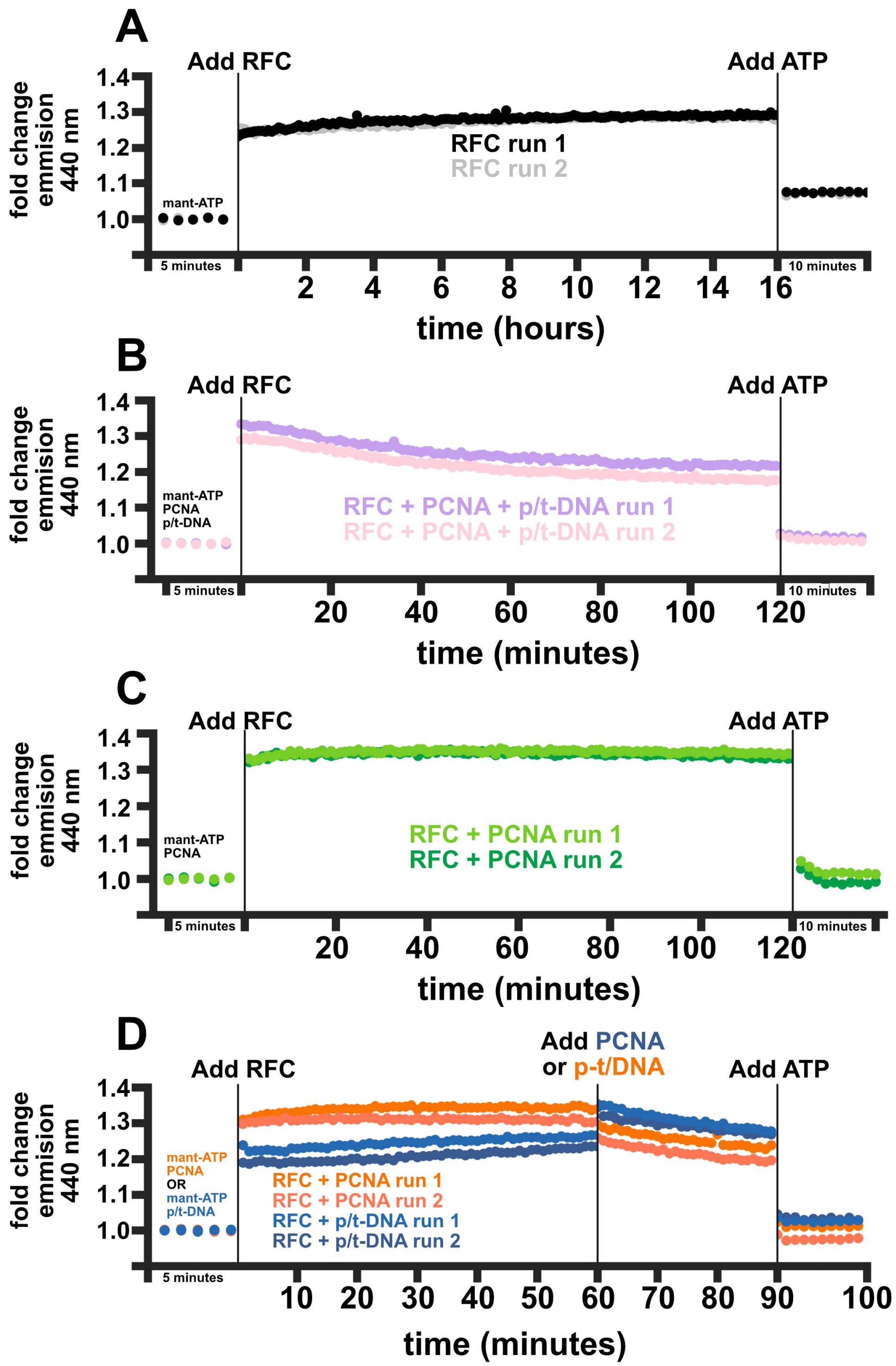


**Supplemental Figure 5S3. MANT-ATP binding kinetics to RFC.** In all cases, the five minutes leading up to addition of RFC is used to set the baseline, and fluorescence is reported as a fold change above this baseline. **A.** RFC on its own. **B.** MANT-ATP was preincubated with PCNA and p/t-DNA before adding RFC. **C.** MANT-ATP was preincubated with PCNA before adding RFC. **D.** MANT-ATP was either preincubated with PCNA or p/t-DNA before adding RFC. After one hour, the other component was added.

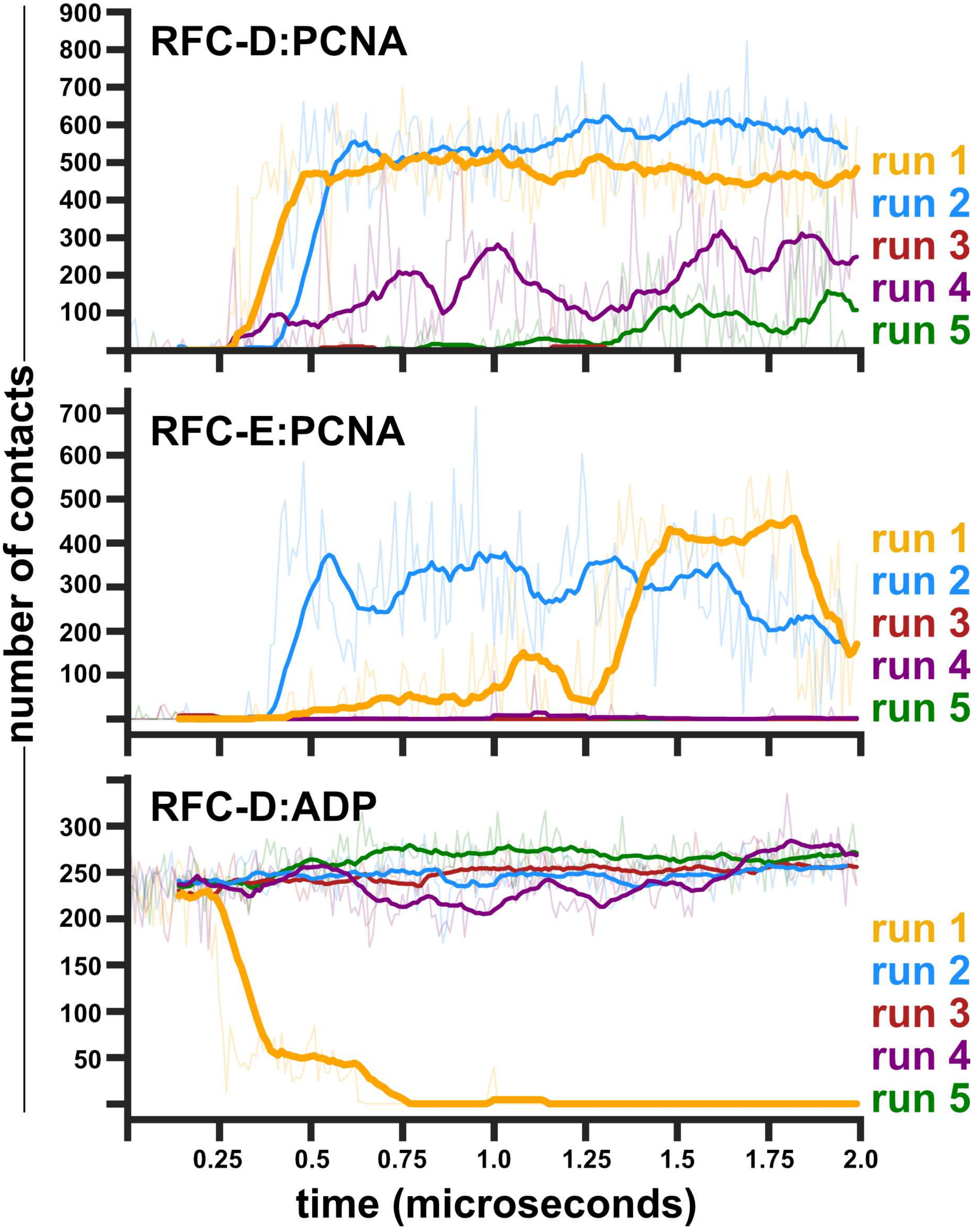


**Supplemental Figure 6S1. Contacts between RFC subunits and PCNA or ADP.** The number of contacts (criterion: heavy-atom within 4.5 Å) between the RFC-D:PCNA, RFC-E:PCNA, and the RFC-D:ADP are shown. Instantaneous values are shown as semi-transparent lines, and a rolling average is shown as thick lines. In run 1, D engages PCNA at ~0.3 μs, while E does not engage until ~1.25 μs. During this time, the interface is pried apart, and ADP is released from the active site. D also engages with PCNA in run 2, but this engagement is concomitant with E engaging PCNA. So, the interface is not pried apart and ADP is not released. D & E only transiently contact PCNA in the other three simulations.
