## Supplementary material for "PCNA is a Nucleotide Exchange Factor for the Clamp Loader ATPase Complex": molecular dynamics input files: README.txt

This folder contains NAMD input files associated with the manuscript
"PCNA is a Nucleotide Exchange Factor for the Clamp Loader ATPase Complex"
by Pajak et al.

The parent folder contains files common to all systems.

toppar contains all force field files for NAMD, including toppar_water_ions_cufix
necessary for applying the CUFIX nonbonded corrections to C36m.

Equilibration is done in three steps, eq.inp followed by eq2.inp and lastly eq3.inp.
Production is performed with the prod.inp file.

Each subfolder contains starting coordinates and topology files for each system simulated.
    adp contains files for ADP-bound RFC
    atp contains files for ATP-bound RFC
    pcna contains files for RFC:PCNA with ADP in the RFC-D subunit

.inp files needed for performing the tauRAMD simulations are found in the adp folder
ADP release from each subunit has a unique .inp files which specifies the starting and
stopping atoms for ADP & the Rossmann fold of the associated subunit. 
To run replicates, one should change ramdSeed to some other value.
